# *Vivaxin* genes encode highly immunogenic non-variant antigens unique to the *Trypanosoma vivax* cell-surface

**DOI:** 10.1101/2022.02.16.480502

**Authors:** Alessandra Romero-Ramirez, Aitor Casas-Sánchez, Delphine Autheman, Craig W. Duffy, Marta M. G. Teixeira, Rosangela Z. Machado, Janine Coombes, Robin J. Flynn, Gavin J. Wright, Andrew P. Jackson

## Abstract

*Trypanosoma vivax* is a unicellular hemoparasite, and a principal cause of animal African trypanosomiasis (AAT), a vector-borne and potentially fatal disease of livestock across sub-Saharan Africa. Previously, we identified diverse *T. vivax*-specific genes that were predicted to encode cell surface proteins. Here, we examine the immune responses of naturally and experimentally infected hosts to many of these unique parasite antigens, to identify immunogens that could become vaccine candidates. Immunoprofiling of host serum showed that one particular family (Fam34) elicits a consistent IgG antibody response. This gene family, which we now call *Vivaxin*, encodes at least 124 transmembrane glycoproteins that display quite distinct expression profiles and patterns of genetic variation. We focused on one gene (viv-*β*8) that is among the most immunogenic and highly expressed but displays minimal polymorphism. VIVβ8 was localized across the cell body and flagellar membrane, suggesting that vivaxin is substantial family of novel surface proteins. Although vaccination of mice with VIVβ8 adjuvanted with Quil-A elicits a strong, balanced immune response and delays parasite proliferation in some animals, ultimately, it does not prevent disease. However, our phylogenetic analysis shows vivaxin includes other antigens shown to induce immunity against *T. vivax*. Thus, the introduction of vivaxin represents an important advance in our understanding of the *T. vivax* cell surface. Besides being a source of proven and promising vaccine antigens, the gene family is clearly an important component of the parasite glycocalyx, with potential to influence the host-parasite interaction.

**Author summary:** Animal African trypanosomiasis (AAT) is an important livestock disease throughout sub-Saharan Africa and beyond. AAT is caused by Trypanosoma vivax, among other species, a unicellular parasite that is spread by biting tsetse flies and multiplies in the bloodstream and other tissues, leading to often fatal neurological conditions if untreated. Although concerted drug treatment and vector eradication programmes have succeeded in controlling Human African trypanosomiasis, AAT continues to adversely affect animal health and impede efficient food production and economic development in many less-developed countries. In this study, we attempted to identify parasite surface proteins that stimulated the strongest immune responses in naturally infected animals, as the basis for a vaccine. We describe the discovery of a new, species-specific protein family in T. vivax, which we call vivaxin. We show that one vivaxin protein (VIVβ8) is surface expressed and retards parasite proliferation when used to immunize mice, but does not prevent infection. However, we also reveal that vivaxin includes another protein previously shown to induce protective immunity (IFX/VIVβ1). Besides its great potential for novel approaches to AAT control, vivaxin is revealed as a significant component of the T. vivax cell surface and may have important, species-specific roles in host interactions.

## Introduction

African trypanosomes *(Trypanosoma subgenus Salivaria)* are unicellular flagellates and obligate hemoparasites. *Trypanosoma vivax* is one of several African trypanosome species that cause the animal African trypanosomiasis (AAT), a vector-borne disease of livestock that is endemic across sub-Saharan Africa, as well as found sporadically in South America [1-2]. Cyclical transmission of T. *vivax* by tsetse flies *(Glossina spp*.), or mechanical transmission by diverse other biting flies, leads to an acute, blood-borne parasitaemia and subsequent chronic phases during which parasites disseminate to various tissues, the central nervous system in particular [1-5]. AAT is a potentially fatal disease characterised by acute inflammatory anaemia and various reproductive, neural and behavioural syndromes during chronic phase [6-7]. The impact of the disease on livestock productivity, and therein the food security and wider socio-economic development of endemic countries, is profound and measured in billions of dollars annually [8]. Thus, AAT is rightly considered one of the greatest challenges to animal health in these regions [9-10].

Strategies to prevent AAT are typically based around vector control, using insecticides, traps or pasture management, in combination with prophylaxis with trypanocidal drugs [11]. However, widespread drug resistance and the on-going cost of maintaining transnational control means that, in an ideal situation, a vaccine is the preferred, sustainable solution [12-13]. African trypanosome infections are, however, far from an ideal target for vaccination for two reasons. First, antigenic variation of the Variant Surface Glycoprotein (VSG) enveloping the trypanosome cell leads to immune evasion and immunization with VSG fails to protect against heterologous challenge [14]. Second, chronic inflammation induced by infection leads to oblation of memory B-cells and immunosuppression [13].

Successful recombinant vaccines exist for other pathogens that are capable of antigenic switching, for example, hemagglutinin of influenza [15], hepatitis C [16] outer surface antigens of Borrelia [17] and the circumsporozoite protein of *Plasmodium falciparum* [18]. These vaccines are based on pathogen surface antigens that elicit dominant immune responses in natural infections. Thus, while antigenic variation of trypanosomes specifically precludes whole-cell vaccine approaches, recombinant vaccines might work if based on non-VSG antigens exposed to the immune system during infections. Yet, most experiments using various conserved and invariant trypanosome proteins [19-21] have not led to robust protective immunity, causing the very plausibility of African trypanosome vaccines to be questioned [22]. Recently, however, in a systematic screen of recombinant subunit vaccines based on T. *vivax* non-VSG surface antigens, we identified a T. *vivax*-specific, invariant flagellum antigen (IFX) that induced long-lasting protection in a mouse model. This immunity was passively transferred with immune serum, and recombinant monoclonal antibodies to IFX could induce sterile protection [23].

In this study, we continue our evaluation of T. *vivax* antigens using a complementary approach, beginning by analysing the natural immune responses to T. *vivax*-specific surface proteins. We previously categorized genes encoding T. vivax-specific, cell-surface proteins that were not VSG (‘TvCSP’) into families, named Fam27 to Fam45 inclusive [24]. We showed that many of these TvCSP families (e.g. Fams 29, 30, 32, 34 and 38) are abundant and preferentially expressed in bloodstream-form parasites [25]. Our aim here is to identify candidates for recombinant vaccine development through four objectives: (1) to assay serum from naturally infected animals using a custom TvCSP peptide array; (2) to produce recombinant protein for immunogenic TvCSP using a mammalian expression system; (3) to confirm the cell-surface localisation of TvCSP using immunofluorescent microscopy; and (4) to vaccinate and challenge with T. *vivax* in a mouse model.

We show that one TvCSP family of 124 paralogous genes encoding putative type-1 transmembrane proteins are especially immunogenic in natural infections, and we name this family vivaxin. Vaccination with *vivaxin* produces a robust, mixed immune response in mice that significantly reduces parasite burden, but without ultimately preventing infection. We show that at least one vivaxin family member is found on the extracellular face of the plasma membrane of T. vivax bloodstream-stage trypomastigotes, and therefore, aside from its utility as a vaccine candidate, *vivaxin* is likely to be an abundant component of the native T. vivax surface coat, alongside VSG.

## Methods

### Ethical approvals

All mouse experiments were performed under UK Home Office governmental regulations (project licence numbers PD3DA8D1F and P98FFE489) and European directive 2010/63/EU. Research was ethically approved by the Sanger Institute Animal Welfare and Ethical Review Board. Mice were maintained under a 12-h light/dark cycle at a temperature of 19–24⍰°C and humidity between 40 and 65%. The mice used in this study were 6–14-week-old male and female Mus musculus strain BALB/c, which were obtained from a breeding colony at the Research Support Facility, Wellcome Sanger Institute.

### Design and production of TvCSP peptide microarray

The array design included 65 different T. *vivax* Y486 antigens (45 representatives of TvCSP multi-copy families and 20 T. *vivax*-specific, single copy genes with predicted cell surface expression), and one neo epitope. The microarrays comprised 600 peptides printed in duplicate, each 15 amino acids long with a peptide-peptide overlap of 14 amino acids, and manufactured by PEPperPRINT (Heidelberg, Germany). Each array included peptides cognate to mouse monoclonal anti-FLAG (M2) (DYKDDDDKAS) and mouse monoclonal influenza hemagglutinin HA (YPYDVPDYAG), displayed on the top left and bottom right respectively, which were used as controls (12 spots each control peptide).

### Infected host serum

Blood serum from cattle known, or suspected, to be infected with T. *vivax* were obtained from Kenya (N = 24), Cameroon (N = 26) and Brazil (N = 4). African samples came from naturally infected animals, while Brazilian serum came from calves experimentally infected with the Brazilian T. vivax Lins strain. Samples were screened with the Very Diag diagnostic test (Ceva-Africa; [26]), which confirmed that they were seropositive for T. *vivax*. Negative controls were provided by serum from UK cattle (N = 4), seronegative by diagnostic test. A further control for cross-reactivity with T. congolense, (commonly co-incident with T. *vivax*), utilised serum from Cameroonian cattle (N = 11) that were seronegative by diagnostic test for T. *vivax*, but seropositive for T. *congolense*.

### Immunoprofiling assay

Fifteen of the 57 positive T. *vivax* samples to be tested in the microarrays were seropositive for T. *vivax* only (i.e. unique infection), while 42 were seropositive for both T. vivax and T. *congolense*. Before applying these to the peptide arrays, one array was pre-stained with an anti-bovine IgG goat secondary antibody (H+L) Cy3 (Jackson ImmunoResearch Laboratories) at a dilution 1:4500 in order to obtain the local background values. Slides were analyzed with an Agilent G2565CA Microarray Scanner (Agilent Technologies, USA) using red (670nm) and green (570nm) channels independently with a resolution of 10um. The images obtained were used to quantify raw, background and foreground fluorescence intensity values for each spot in the array using the PEPSlide Analyzer software (Sicasys Software GmbH, Heidelberg, Germany).

### Immunoprofiling analysis

A cut-off threshold was defined according to Valentini *et al*. and applied to the raw intensity values [27]. The limma R package from Bioconductor [28] was used to identify top immunogenic peptides in livestock serum samples. The data import was extracted directly from the Genepix files (.gpr) produced by the PepSlide Analyzer, using only the green channel intensity data. The “normexp” method was selected for background and the normalization between arrays achieved with vsn [29]. A filtering step was performed removing control peptides (HA and FLAG) from each array and the intensity values from spots in duplicate were averaged. Multiple statistic tests for differential expression were calculated (p-value < 0.05) selecting Benjamini and Hochberg’s method for the false discovery rate (log2fold-change (FC) >2) [30]. The peptides were ranked based on their p-values as well as their fold-change to determine which peptides were bound by serum antibodies significantly more than background.

### Phylogenetic analysis

All full-length vivaxin gene sequences (n = 81) were extracted from the T. *vivax* Y486 reference genome sequence. Amino acid sequences were aligned using Clustalx [31] and then back-translated and manually checked using Bioedit [32], producing a 663 nucleotide codon alignment (221 amino acids). Phylogenies were estimated for both codon and amino acid alignments using Maximum likelihood and Bayesian inference. Maximum likelihood trees were estimated using Phyml [33] with automatic model selection by SMS [34], according to the Akaike Information Criterion. The optimal models were GTR+Γ (α = 3.677) and JTT+Γ (α = 5.568) for codon and protein alignments respectively. Topological robustness was measured using an approximate log-likelihood ratio (aLRT) branch test, as well as 100 non-parametric bootstrap replicates. Raxml [35] was also used to estimate bootstrapped maximum likelihood trees, using unpartitioned GTR+FU+Γ (α = 3.134) and LG+Γ (α = 3.847) models for codon and protein alignments respectively. Bayesian phylogenies were estimated from the same alignments using Phylobayes [36], employing four Markov chains in parallel and a CAT model with rate heterogeneity. A single, divergent sequence (TvY486_0024510) was designated as outgroup because it branches close to the mid-point in all analyses.

### Recombinant protein expression

Protein sequences encoding the extracellular domain and lacking their signal peptide, were codon optimized for expression in human cells and made by gene synthesis (GeneartAG, Germany and Twist Bioscience, USA). The sequences were flanked by unique NotI and AscI restriction enzyme sites and cloned into a pTT3-based mammalian expression vector 26 between an N-terminal signal peptide to direct protein secretion and a C-terminal tag that included a protein sequence that could be enzymatically biotinylated by the BirA protein-biotin ligase 32 and a 6-his tag for purification. The ectodomains were expressed as soluble recombinant proteins in HEK293 cells as described [37-38]. To prepare purified proteins for immunisation, between 50 and 1.2L (depending on the level at which the protein was expressed) of spent culture media containing the secreted ectodomain was harvested from transfected cells, filtered and purified by Ni2+ immobilised metal ion affinity chromatography using HisTRAP columns using an AKTAPure instrument (GEHealthcare, UK). Proteins were eluted in 400mM imidazole as described [39] and extensively dialysed into HBS before quantified by spectrophotometry at 280nm. Protein purity was determined by resolving one to two micrograms of purified protein by SDS-PAGE using NuPAGE 4–12% Bis Tris precast gels (ThermoFisher) for 50 minutes at 200V. Where reducing conditions were required, NuPAGE reducing agent and anti-oxidant (Invitrogen) were added to the sample and the running buffer, respectively. The gels were stained with InstantBlue (Expedeon) and imaged using a c600 Ultimate Western System (Azure biosystems). Purified proteins were aliquoted and stored frozen at -20°C until use.

### Cellular localization

Cellular localization of VIVβ8 (‘antigen-4’) in T. *vivax* bloodstream-forms was determined by indirect immunofluorescence. T. *vivax* bloodstream-forms were isolated as described previously [23], adjusted to 2.5×10^6^ cells/ml in PBS+20mM glucose, transferred to poly-L-lysine slides for 10min and fixed either in 4% paraformaldehyde (PFA) or 4% PFA supplemented with 0.25% glutaraldehyde for 30min at RT. Cells were washed with PBS and blocked with blocking buffer (PBS+1% BSA) for 1h at RT. Either pooled anti-VIVβ8 post-immune mouse sera or purified rabbit anti-VIVβ8 IgGs was used as primary antibody (1:1,000 dilution) in blocking buffer and incubated overnight at 4°C. After washing, cells were incubated for 1h at RT with goat anti-mouse IgG conjugated with Alexa Fluor-555 (Abcam, UK) (1:500 dilution in blocking buffer). Cells were incubated in 500 ng/ml DAPI (Invitrogen, USA), and/or 1:100 mCLING unspecific staining (Synaptic Systems), washed and mounted in Slow Fade diamond antifade mounting oil. Cells were imaged using a LSM-800 confocal laser scanning microscope (Zeiss). Images were processed using Zen 3.1 (Zeiss), ImageJ [40]. 3D renders were generated from z-tacks using ImarisViewer 9.5.1 (Imaris).

A polyclonal antibody against recombinant VIVβ8 was raised in rabbits (BioServUK, Sheffield, UK). Briefly, two rabbits were vaccinated by subcutaneous injection, receiving five injections of 0.15mg VIVβ8 antigen diluted in sterile PBS and co-administrated with Freund’s adjuvant every two weeks (0.75mg total immunization). IgG antibodies were purified by affinity chromatography with a protein A column from antisera collected two weeks after the last boost. The final concentration of the rabbit purified antibody was 5mg/ml.

### Live immunostaining

Bloodstream-form T. *vivax* were isolated from infected mice as previously described and incubated with primary anti-VIVβ8 (purified rabbit polyclonal, 1:200 dilution) in blocking solution (1% BSA) for 30 minutes at either 4°C or RT. Cells were washed in PBS 20mM glucose by centrifugation and incubated with secondary Alexa Fluor goat anti-rabbit IgG 555 conjugated in blocking solution for 30 minutes at either 4°C or RT. After washing as described above, all cells were fixed in 4% PFA at for 30 minutes at RT. Cells were then incubated with 500 ng/mL DAPI DNA counterstain, mCLING unspecific staining (1:100 dilution) and/or 5 mg/mL FITC-conjugated ConA for 15 minutes at RT. After washing, cells were mounted in SlowFade diamond mounting oil.

### Western Blotting

A Western blot was performed to confirm the cellular location of VIVβ8 antigen in natural settings. Three gels were run each one to be incubated with a different secondary antibody. Samples were as follows: 1:250 pool of sera from naturally infected cattle from Malawi (n=3), Cameroon (n=4) and Kenya (n=4) and the elution from FTA cards containing plasma from experimentally infected cattle from Brazil (n=6) and a pool of sera from uninfected British cattle (n=4) used as negative control. In addition, 1μg VIVβ8 and 1μg VIVβ14 biotinylated proteins and 1.67×10^6^ lysed parasites of *T. brucei* and *T. vivax*, respectively, were used as controls. The samples were mixed with loading buffer (Laemmli buffer, 2X concentrate) (70), heated for 5min at 95°C and run in an 10% SDS/polyacrylamide gel electrophoresis. The proteins were transferred to polyvinylidene difluoride membrane (Milipore, Billerica, MA, USA) using a semi-dry chamber (Trans blot SD semi-dry transfer cell, Biorad) and blocked with 4% semi-skimmed milk in PBS (blocking buffer) overnight at 4°C. The membranes were washed 3 times with PBS+0.05% Tween 20 and probed with either anti-VIVβ8 or anti-VIVβ14 polyclonal antibodies 1:500 or anti-PFR antibody 1: 1,000 in blocking buffer for 2h at RT. Membranes were washed as before and incubated with goat anti-rabbit IgG HRP-conjugate in blocking buffer for 2h at RT. The membranes were washed as before, revealed (Clarity Western ECL Substrate, BioRad) and bands were visualized with a ChemiDoc MP Imaging system (BioRad).

### Vaccine preparation

VIVβ11, VIVβ14, VIVβ20 and VIVβ8 recombinant proteins were combined independently with one of the three adjuvants to analyze the potential different types of immune responses. The vaccine formulation was prepared by combining 20μg purified antigen with either 100μg Alum (InvivoGen), Montanide W/O/W ISA 201 VG (Sappec, France) or 15μg saponin Quil-A (Invivogen, USA), respectively. Moreover, an additional vaccine was formulated with 50μg VIVβ8 and 15μg Quil-A. Control animals were immunized with the adjuvants only using the same concentration as the antigen-vaccinated groups.

### Mouse immunization and challenge with T. vivax

Male mice were distributed in groups (n=3) as follows for the immunization: four groups were immunized with Alum in combination with each antigen, (i.e. VIVβ*/A); and four groups with each antigen co-administrated with Montanide, (i.e. VIVβ*/M). There was one control group each for VIVβ*/A or VIVβ*/M-vaccinated group. In addition, female mice were randomly distributed in five groups (n=8), four of them immunized with Quil-A plus each antigen (i.e. VIVβ*/Q) and one as control group immunized with adjuvant only. Mice from all groups were immunized on days 0, 14 and 28 subcutaneously in two injection sites (100μl/injection). Animals from the VIV*/A and VIV*/M groups were euthanized two weeks after the third immunization (day 42), since it was clear that Quil-A was the preferred adjuvant. The VIVβ*/Q rested for 14 prior to challenge; at day 42, they were infected intraperitoneally with 10^3^ bioluminescent, bloodstream-form *T. vivax* parasites (Y486 strain). The parasites were obtained at day 7 post infection (dpi) from previous serial passages in mice. Briefly, 10μl whole blood was collected and diluted 1:50 with PBS+ 5% D-glucose+10% heparin. The parasite concentration was adjusted to 10^2^ parasites/200μl/mouse. After challenge, the animals were monitored daily and quantification of *T. vivax* infection was measured by bioluminescent in vivo imaging. Subsequently, a second challenge was conducted to confirm the results; two groups of mice (n = 15) were immunized following the same schedule as before with 50μg VIVβ8 + 15μg Quil-A and adjuvant only respectively, prior to challenge on day 74.

### In vivo imaging

Animals were injected daily starting at 5dpi and 6dpi for the first and second challenge respectively with luciferase substrate D-luciferin (potassium salt, Source BioScience, UK) diluted in sterile PBS for in vivo imaging and data acquisition. Mice were injected intraperitoneally with 200 μl luciferin solution at a dose of 200mg/kg per mouse 10 minutes before data acquisition. Animals were anaesthetized using an oxygen-filled induction chamber with 3% isoflurane and bioluminescence was measured using the in vivo imaging system IVIS (IVIS Spectrum Imaging System, Perkin Elmer). Mice were whole-body imaged in dorsal position and the signal intensity was obtained from luciferase expressed in *T. vivax*. The photon emission was captured with charge coupled device (CCD) camera and quantified using Living Image Software (Xenogen Corporation, Almeda, California) The data were expressed as total photon flux (photons/second).

### Serum collection

Blood was collected from the tail vein of each animal at day 0 (pre-immune sera), day 42 (post-immune sera for alum and montanide treatment groups) and day 50 (post-immune sera for VIV*/Q, VIVβ8-B and VIVβ8-C groups). Sera were isolated from blood by centrifuging the samples for 10min x 3,000rpm and the supernatant was stored at -20 °C until used for antibody titration. Spleens were aseptically removed from the VIV*/A and VIV*/M groups 42dpi and from challenged groups of the 1^st^ CH at 50dpi and 2^nd^ CH at 83dpi. Spleen tissue was used for *in vitro* antigen stimulation in order to quantify cytokine expression.

### In vitro antigen stimulation and cytokine measurement

Splenocytes were isolated by collecting spleens individually in tubes containing 3ml sterile PBS. The cells were separated with a 70μm cell strainer attached to a 50ml tube and pressed using a syringe plunger end. The samples were centrifuged at 800g for 3min at RT. Erythrocytes were lysed by adding 2ml ACK lysis buffer (0.15M NH_4_Cl, 10mM KHCO_3_, 0.1mM EDTA, pH 7.5) to the pellet and incubated for 5min. Lysis buffer was neutralized by adding 30ml of complete media (RPMI 1640 (Sigma Aldrich, Germany) supplemented with 10% heat inactivated foetal calf serum (FCS; Sigma Aldrich, Germany), 100 U/ml penicillin and 100U/ml streptomycin) and centrifuged as before. Cell density was adjusted to 5×10^6^ cells/ml per spleen in complete medium and cultured in 48-well flat-bottom tissue culture plates (Starlab, UK) by seeding 200μl/well each suspension in triplicate. Splenocytes were stimulated with 10μg/ml each antigen diluted in complete medium for 72h at 37°C with 70% humidity and 5% CO_2_. Likewise, cells were also incubated with 10μg/ml Concanavalin A (ConA) or complete medium only as positive and negative controls, respectively. Culture supernatants were harvested after 72h and centrifuged at 2000g for 5min at RT to remove remaining cells. The supernatant was collected and used for the quantification of interferon gamma (IFN-γ), tumour necrosis factor (TNF-α), interleukin-10 (IL-10) and interleukin-4 (IL-4) levels by sandwich ELISA kits (ThermoFisher Scientific). The measurement from unstimulated splenocytes (incubated with medium only) was subtracted from the antigen stimulated cultures with each adjuvant treatment.

### IgG-specific antibody response in mice and natural infections

To identify the presence of specific antibodies in mice sera against the antigens, a titration of IgG1 and IgG2a isotypes was performed by indirect ELISA. Briefly, 96-well streptavidin-coated plates were incubated with each antigen for 1h at RT with 1:250 VIVβ11 and 1:50 VIVβ14/Q, VIVβ20/Q and VIVβ8/Q diluted in reagent diluent (PBS pH 7.4, 0.5% BSA). The plates were washed three times with PBS-Tween20 0.05% and two-fold serial dilutions of each serum diluted in reagent diluent were performed, added to each well and incubated for 1h at RT. Plates were washed as before and 100μl/well rabbit anti-mouse IgG1 or IgG2a conjugated to HRP (Sigma-Aldrich, Germany) diluted to 1: 50,000 and 1: 25,000, respectively, were added to the plates and incubated as before. After washing, 100μl/well of 3,3’,5,5’-tetramethylbenzidine (TMB, Sigma-Aldrich, Germany) was incubated for 5 minutes at RT in the dark. The reaction was stopped by adding 50μl/well 0.5M HCl and the absorbance was read at 450nm using a Magellan Infinite F50 microplate reader (Tecan, Switzerland).

The isotype profile against each antigen was also analysed in samples from naturally and experimentally infected cattle. The ELISA protocol used was the same as above performing two-fold serial dilutions in samples from experimental and natural infections. Bound IgG1 and IgG2 antibodies were detected by adding 100μl/well sheep anti-bovine IgG1 or IgG2 HRP (Bio-Rad, USA) at 1:5000 and 1:2500 respectively.

## Results

### Immunoprofiling of naturally infected livestock serum identifies consistent T. vivax-specific antigens

The immuno-reactivity of serum from animals seropositive for *T. vivax* to diverse *T. vivax*-specific antigens was examined using a custom peptide microarray of 66 putative cell surface proteins (S1 Fig). Serum taken from natural bovine infections in Kenya and Cameroon, as well as experimental ovine infections with Brazilian *T. vivax* strains, gave a consistent response, shown in Fig 1. It is immediately clear from the arrays that infected sera produce a consistent response, with spots in the two top-most rows of the array responding strongly, while corresponding positions in negative controls (both UK cattle that were *T. vivax* seronegative and Kenyan cattle that were seropositive for *T. congolense* only) lack this response (Fig 1A). The pattern remains after normalization for variation within and between arrays, with Kenyan, Cameroonian and Brazilian samples displaying a spike in intensity for these spots that is missing in negative controls (Fig 1B). Of the 60 spots in the top row of Figure 1, 51 correspond to Fam34 (see S1 Fig), previously described as a family of putative transmembrane proteins and highly abundant in bloodstream-stage mouse infections (Jackson et al. 2015).

**Figure 1.**
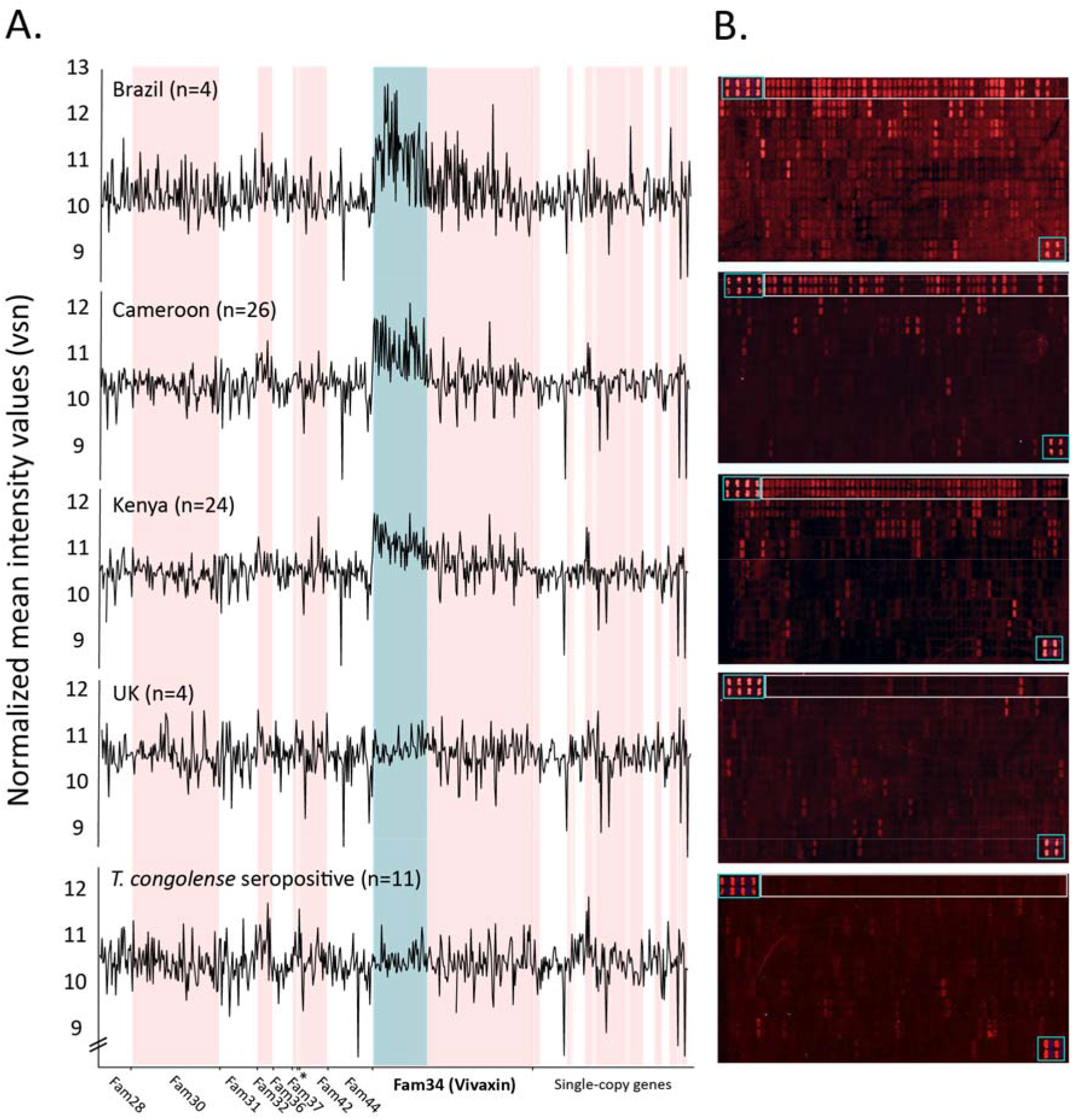
Immuno-profiling of host serum with a Trypanosoma vivax-specific antigen array. **(A)** Fluorescence observed after application of host sera to a customised array of 600 peptides representing 66 T. vivax-specific antigens. From top to bottom, the panels show responses to serum from three locations (Brazil, Cameroon and Kenya respectively), in addition to two negative controls (serum from UK cattle and from Cameroonian cattle that tested *T. vivax*-negative *but T. congolense*-positive). Control peptides are shown in boxes at top left and bottom right of each array. The top row of the array (boxed) contains peptides exclusively derived from vivaxin proteins. **(B)** Normalized intensity values of immuno-fluorescent responses in (A) are plotted for all array peptides, arranged by gene family. Individual gene families are indicated by alternating white and pink shading. Strongly-responding peptides belonging to vivaxin (Fam34) are indicated by blue shading.

Table 1 describes the spots with the strongest response intensity values (highest 10% values) and shows which spots were statistically significant, i.e. were significantly greater than background fluorescence determined by the negative control (see Methods). Of these 60 strongly responding spots, 45 relate to peptides derived from Fam34 proteins. Moreover, these are particular Fam34 proteins. 29/45 relate to a single family member (hereafter ‘antigen 1’), nine more relate to a second protein (‘antigen-2’), three relate to ‘antigen-3’ and two to ‘antigen-4’. Peptides from antigens 1 and 2 have the highest maximum fold-change in normalized fluorescence intensity relative to the negative control, e.g. peptide 45 (4.53) and peptide 35 (2.38), and, after adjustment for multiple tests, four peptides belonging to antigen-1 (shown in Table 1), and two more from antigen-2, have response values that exceed the significance threshold (p < 0.05).

**Table 1.**
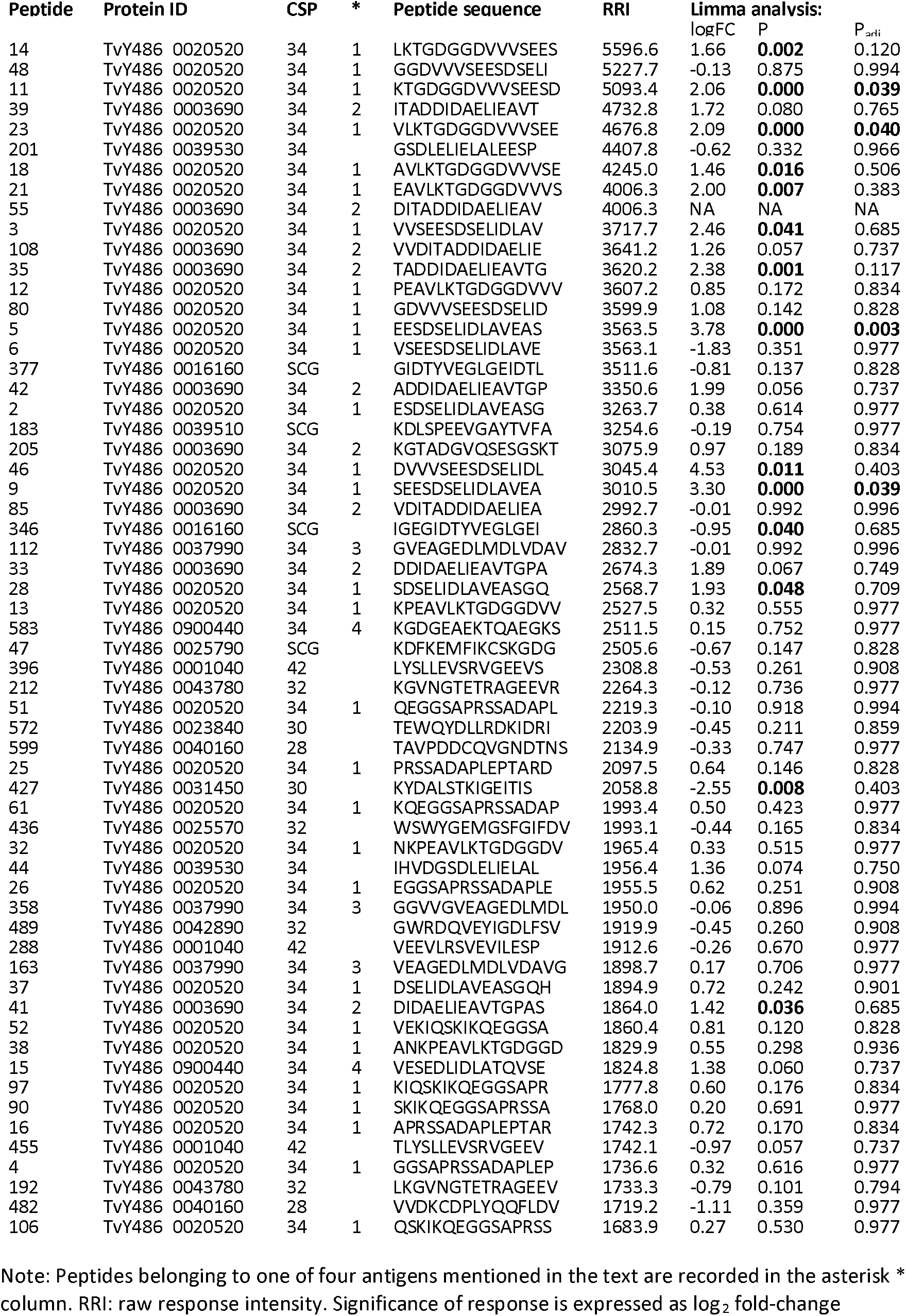
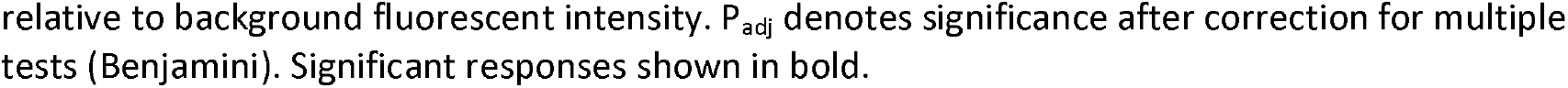
Parasite peptides displaying the strongest responses to host antibodies, (highest 10% when ranked by raw response intensity), in an immuno-profiling assay of *T. vivax*-infected serum.

Thus, we may conclude that Fam34, and in particular ‘antigen-1’ and ‘antigen-2’, are principally responsible for the peak clearly visible in Figure 1A. Given the pre-eminence of Fam34 proteins as consistent and robust antigens in natural infections, we focused our search for vaccine targets on this gene family, which we now rename *vivaxin*.

### Vivaxin is a species-specific gene family encoding type-1 transmembrane proteins that do not display antigenic variation

Analysis of vivaxin amino acid sequences with BLAST returns no matches outside of *T. vivax* itself, and more sensitive comparison of protein secondary structural similarity using HMMER also fails to detect homologs beyond T. vivax; this confirms that the family is species-specific. Comparison with the *T. vivax* Y486 reference genome (TritrypDB release 46) using BLASTp returns 50 gene sequences, while a further 74 homologs are detected by HMMER, which means that vivaxin is the largest T. vivax cell-surface gene family after VSG (Jackson et al. 2012, 2013). These gene sequences range from 1050-1900 bp in length when complete; 43/124 sequences are curtailed by sequence gaps in the current assembly. Only six sequences are predicted to contain internal stop codons, suggesting that pseudogenes are rare. We observe that almost all BLAST matches relate to sub-telomeric loci, (i.e., outside of regular core polycistrons). Previously, in silico predictions based on amino acid sequences indicated that all Fam34 genes encode a type-1 transmembrane protein with a predicted signal peptide and a single hydrophobic domain 15 amino acids from the C-terminus, orientated such that the protein is largely extracellular [24]. We carried out further analysis of antigens 1-4 with PredictProtein [41] and ModPred [42], shown in S2 Fig, that confirm this topology and suggest that the extracellular portion of vivaxin is both N-and O-glycosylated at multiple sites.

We estimated a Maximum Likelihood phylogeny for 81 full-length vivaxin genes from an alignment of a 221-amino acid conserved region (see Methods). Fig 2A shows that vivaxin sequences group into three robust clades, which we term the α (41 genes), β (34 genes) and γ (5 genes) subfamilies. One additional sequence belonging to TvY486_0024510 diverges close to the mid-point of the tree and was designated as an outgroup. The subfamilies consistently differ in length due to the N-terminal (extracellular) domain of vivaxin-α proteins being □ 200 amino acids longer than vivaxin-β (Fig 2B). For each gene, the proportion of its protein sequence predicted in silico to be B-cell epitope is shown in Fig 2C; on average, 36.1% of a vivaxin protein sequence is predicted to be immunogenic, rising to almost 60% in some cases.

**Figure 2.**
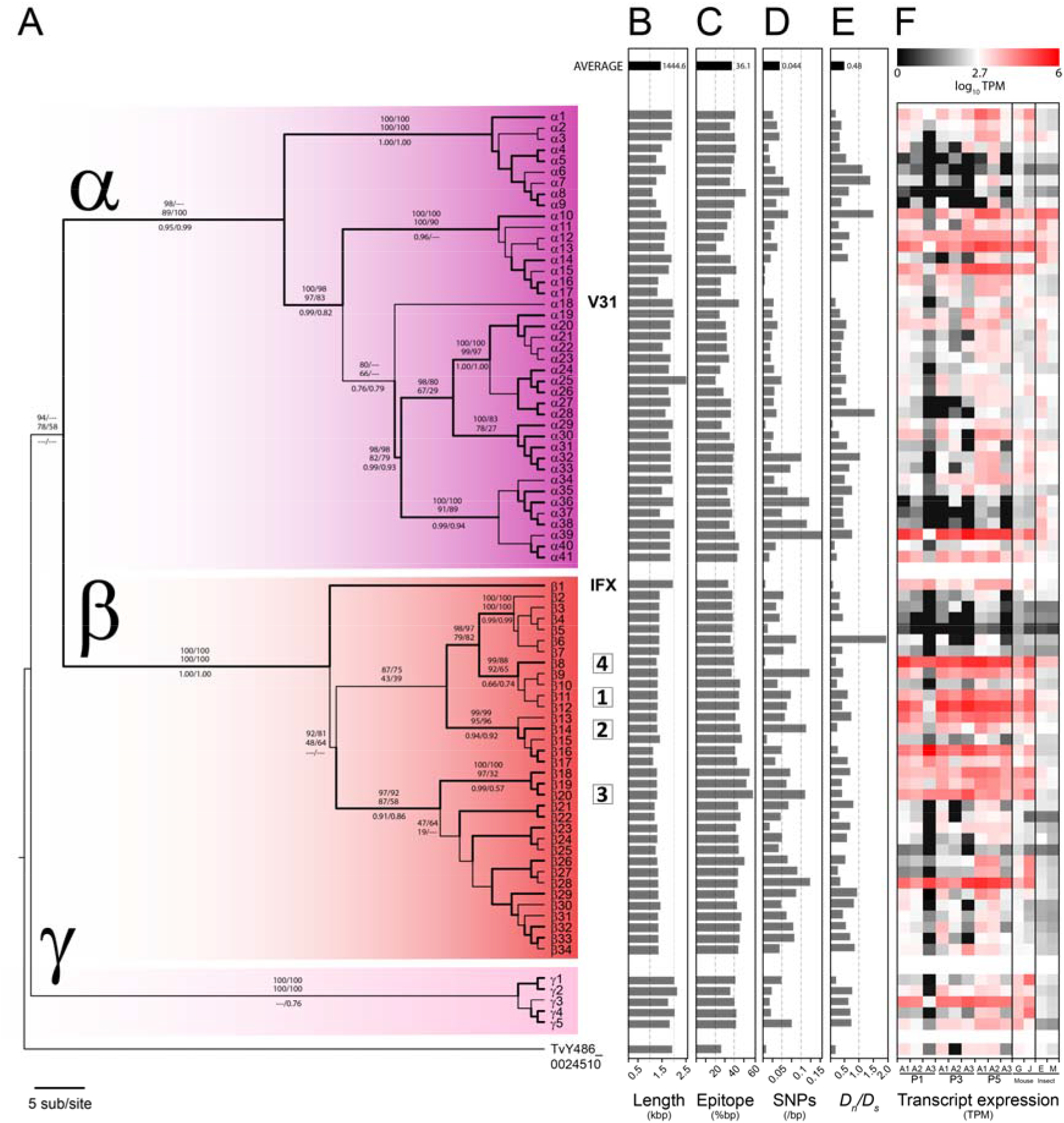
Vivaxin gene family phylogeny and molecular evolution. **(A)** Maximum likelihood phylogeny of vivaxin genes (n = 81) in the *T. vivax* Y486 reference genome estimated with a GTR + Γ substitution model (α = 3.677), and divided into three principal sub-families, labelled α, β and γ. The tree is rooted with a divergent sequence (TvY486_0024510) that approximates to the mid-point. Topological robustness is measured by the approximate log-likelihood ratio (aLRT), and indicated by branch thickness. Thick branches subtend nodes with aLRT values > 0.9. Robustness measures are given for major internal nodes: maximum likelihood bootstrap values (> 75) for nucleotide/protein alignments (upper, above branches), neighbour-joining bootstrap values (> 75) for nucleotide/protein alignments (lower, above branches), and Bayesian posterior probabilities (> 0.5) for nucleotide/protein alignments (below branch). Gene names are arbitrary to protect the identity of possible vaccine candidates. The position of four expressed antigens in this study is indicated by black arrows. **(B)** Gene length mapped to tree topology. **(C)** Total length of predicted B-cell epitopes as a proportion of gene length, as inferred by Bepipred linear prediction 2.0 [45]. **(D)** Single nucleotide polymorphisms (SNP) across a panel of 25 T. vivax strain genomes, as a proportion of gene length. (E) Ratio of non-synonymous to synonymous substitutions (D_n_D_s_) inferred from published SNP data [43]. (F) Heat maps showing vivaxin gene expression profiles from published T. vivax transcriptomes. The first nine columns show relative transcript abundance during an experimental infection in goats. Three peaks in parasitaemia are shown (first, third and fifth respectively; see [43]), with three replicates for each (A1-A3). Columns 10 and 11 show relative transcript abundance in bloodstream-stage infections in mice using different T. vivax strains, LIEM-176 [44] and IL1392 [25], respectively. Columns 12 and 13 show transcript abundance in batch transcriptomes of in vitro cultured T. vivax insect-stages, i.e. epimastigotes (E) and metacyclic-forms (M) respectively [25].

SNPs were identified for each gene using GATK based on variations among diverse clinical *T. vivax* genome sequences we published previously (Fig 2D). These show that, far from being uniformly polymorphic, the population history of vivaxin genes is extremely variable, with some loci being well conserved, indeed almost invariant, across populations. Note that the genes encoding antigens 1-4, (now known as viv-*β*11, -*β*14, -*β*20 and -*β*8 respectively), are all among the least polymorphic paralogs. Some loci, for example viv-*α*15, -*α*29, as well as the gene encoding antigen-4 (viv-*β*8), are predicted to be under highly stringent purifying selection (d_n_/d_s_ ≈ 0; Fig 2E), while others such as viv-*β*9 and these genes encoding ‘antigen-2’ (viv-*β*14) and ‘antigen-3’ (viv-*β*20) appear to be evolving under positive selection (dn/d_s_ > 1). This indicates, first, that there is consistent variation across the family in gene function, with some loci being essential while others are redundant, and second, that this variation has been stable throughout the species history, and not subject to assortment or homogenisation by recombination.

### Vivaxin loci display conserved variation in gene expression profiles

We examined RNAseq data from multiple previous experiments to consider functional variation among vivaxin genes. Fig 2F shows transcript abundance at sequential points of an experimental goat infection ([43]; first nine columns), also in two separate experimental infections in mice ([25,44]; columns 10 and 11) and, finally, in epimastigote (E) and metacyclic (M) parasite stages ([25]; columns 12 and 13 respectively). Most vivaxin genes are expressed weakly in fly stages, confirming that this is predominantly a bloodstream-stage family; although there are exceptions (see viv-*α*36 and α38). Some genes are expressed rarely in all situations, such as viv-*α*6, -*α*8, and -*β*3-6, indicating that they may be non-functional (in the case of viv-*α*6 and -*α*8, these genes do indeed have internal stop codons). Conversely, genes such as viv-*α*10, *α*12, *α*39 and *β*11-12 are expressed constitutively. These genes remain abundant across sequential peaks of bloodstream infections, and across life stages, and indeed, across experiments using different parasite strains and hosts. This clearly indicates that, like many multi-copy surface antigen gene families, expression levels vary markedly among vivaxin genes, but, unlike other gene families, these differences are not dynamic. There is a cohort of vivaxin loci that are routinely active and orders of magnitude more abundant than their congeners. Since many vivaxin genes are expressed constantly and simultaneously, this indicates that they are not variant antigens.

Most interestingly, Figure 2F shows that the genes encoding antigens 1-4 are among the most abundant vivaxin transcripts in all conditions, perhaps explaining why they elicit some of the strongest immune responses. In particular, *viv*-*β*8, which encodes antigen-4, is perhaps the most abundant form of vivaxin, often two orders of magnitude more abundant than most other loci. Taking the results together, we see that the most immunogenic Fam34 proteins (‘antigens 1-4’) are also among the most abundant vivaxin transcripts and among the most evolutionarily conserved vivaxin genes. Our decision to focus on antigens 1-4 as potential subunit vaccines was based on the balance of immunoprofiling, gene expression and polymorphism data.

### Recombinant expression of four β-vivaxin proteins

The four vivaxin proteins were expressed using an AVEXIS assay [37]; in each case, the entire ectodomain was expressed in HEK293-6E mammalian cells as a soluble recombinant protein. The expression of these four *T. vivax* recombinant, non bio-tagged proteins was quantified by western blot and extinction coefficient calculation. The blot performed showed the presence of a prominent band with apparent molecular weight of 50kDa for each recombinant protein (S3 Fig). The antigens have a predicted molecular weight of 34-39KDa based on amino acid sequence alone, i.e. before glycosylation. Based on the extinction coefficient calculation, the purified proteins had a concentration of 4.3μg/ml (VIVβ11), 5.1μg/ml (VIVβ14), 9.8μg/ml (VIVβ20) and 2.5 μg/ml (VIVβ8). Note that weaker, higher molecular weight bands were also observed for all antigens possibly to non-specific binding, the result of post-translational modifications or protein aggregation. These results confirm the abundant expression of authentic, recombinant vivaxin, purified in soluble form, suitable for vaccination.

### Immunofluorescent microscopy localizes viv-*β8* to the whole-cell surface

As yet, the cell-surface position of vivaxin is predicted based on amino acid sequence but not proven. Given the particularly abundant expression and sequence conservation of viv-*β*8 (i.e. antigen-4), we chose to localize this protein by immunostaining *T. vivax* bloodstream forms with anti-VIVβ8 polyclonal antibodies (Fig 3A). When bloodstream form cells were isolated from infected mouse blood; positive stain was associated with the margins of the cell body and flagellum, indicating a specific association with the whole cell surface (Fig 3A, second row). Nonetheless, some cells also showed evidence for intracellular staining, with a noticeable posterior-to-anterior gradient in signal, and a concentrated intensity between the nucleus and kinetoplast (Fig 3A, third row). These observations are not contradictory; endosomes servicing the secretory pathway are known to accumulate at the posterior end of the cell [46], making the intracellular localisation of VIVβ8 consistent with it being trafficked to the cell surface. In fact, surface localisation was confirmed by confocal 3D reconstructions of bloodstream-form cells stained with anti-VIVβ8 antibodies, in where orthogonal views show ring-shaped signal representing the cell edges (Fig 3B).

**Figure 3.**
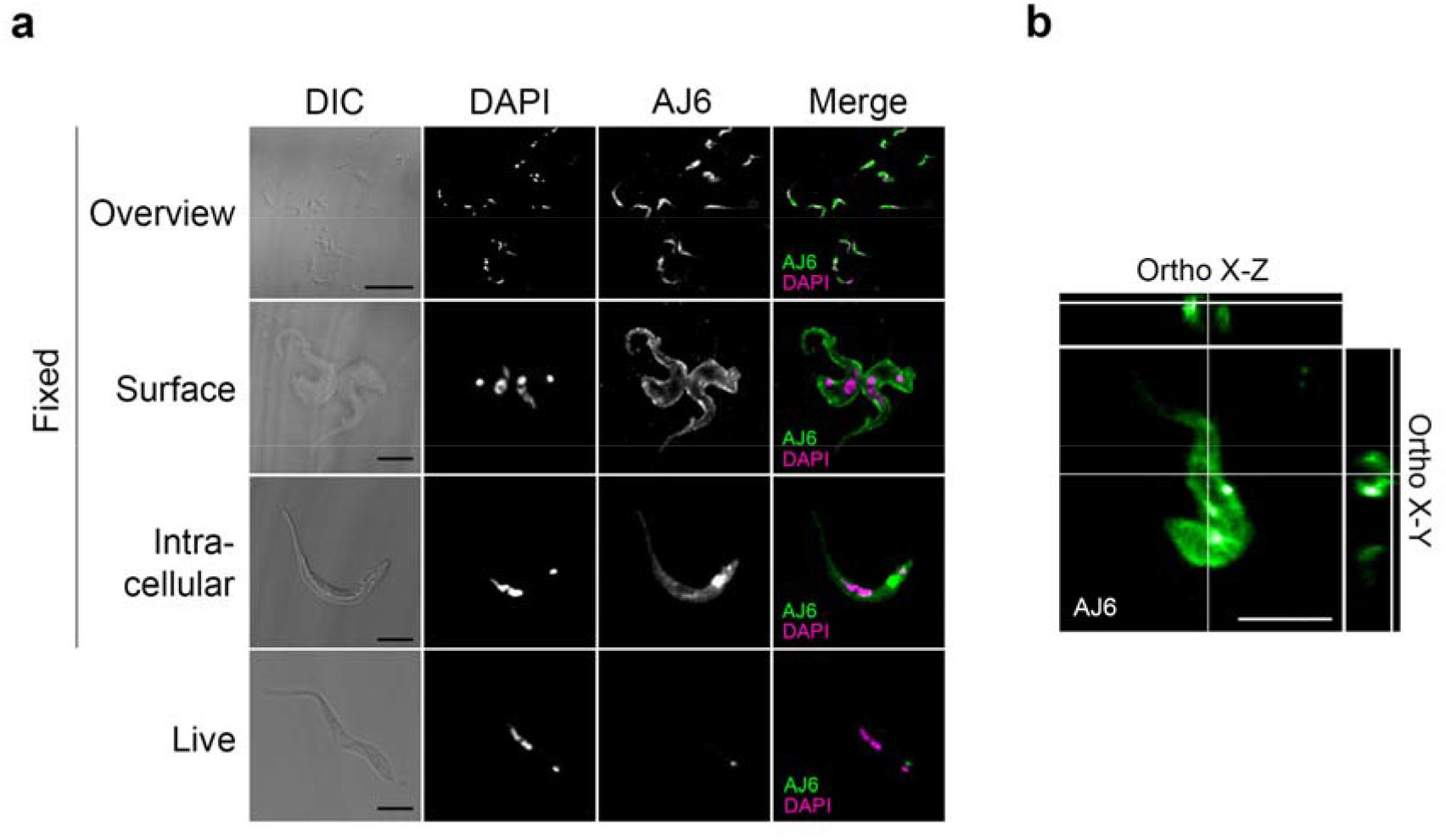
Cellular localization of VIVβ8 protein. **(A)** Representative images of immunofluorescence assays on T. vivax bloodstream forms fixed in either 4% paraformaldehyde (PFA) or PFA supplemented with 0.25% glutaraldehyde (glut), or on live cells. Differential increased contrast (DIC); DAPI DNA counterstain; VIVβ8 (secondary antibody AF555-conjugated) and merged channels. PFA-fixed cells exhibit three main staining patterns; cell surface (Surface), increasing gradient from anterior to posterior end (Gradient), and intracellular mainly (Intracellular). Glut-fixed cells show no antibody binding and live cells only show antibody recognition at the flagellar pocket (see Figure IF_2 for further evidence). Scale bars; 5 μm. **(B)** 3D z-stack reconstructions of T. vivax cells and corresponding orthogonal (X-Z and X-Y) views from the stacks. Orthogonal views note surface localization (circular edges) of VIVβ8 in *T. vivax*.

To corroborate this localisation further, the post-immune sera of mice and rabbits immunised with recombinant VIVβ8 (residue-residue) was used to stain PFA-fixed bloodstream-stage cells (Fig 4). In both cases, post-immune serum reacted strongly with the entire cell surface and flagellum, resembling the localization found using the polyclonal antibody.

**Figure 4.**
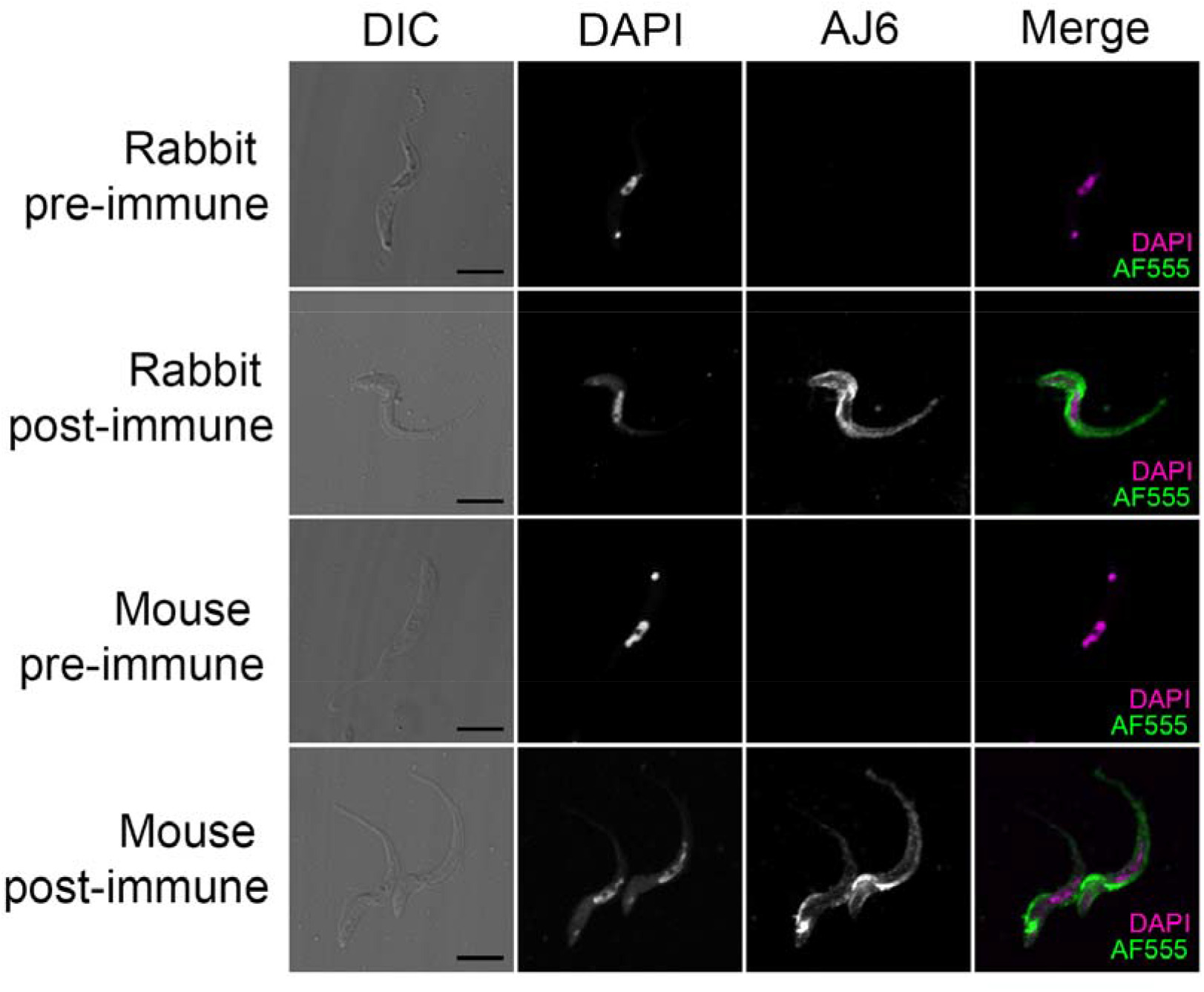
VIVβ8 immunostaining controls. Representative images of *T. vivax* bloodstream-form, PFA-fixed cells either probed with pre-immune (pre) or post-immunization (post) antisera from either rabbit or mouse hosts vaccinated with VIVβ8. Only post-immunisation antisera display antibody binding. Scale bars; 5 μm.

To assess the accessibility of vivaxin epitopes in a native setting, we performed immunostaining on live cells at room temperature (RT) and at 4°C to arrest cell endocytosis (Fig 5A). Unlike in fixed cells, those probed at RT were not stained. However, 4°C-incubated cells localised anti-VIVβ8 exclusively to the flagellar pocket, confirmed by its position next to the kDNA in 3D reconstructions (Fig 5B). As in a previous study [47], we interpret this as evidence for endocytosis. At RT, antibody-bound VIVβ8 is rapidly cleared, but at 4°C, the antibody is not removed and accumulates where VIVβ8 epitopes are exposed.

**Figure 5.**
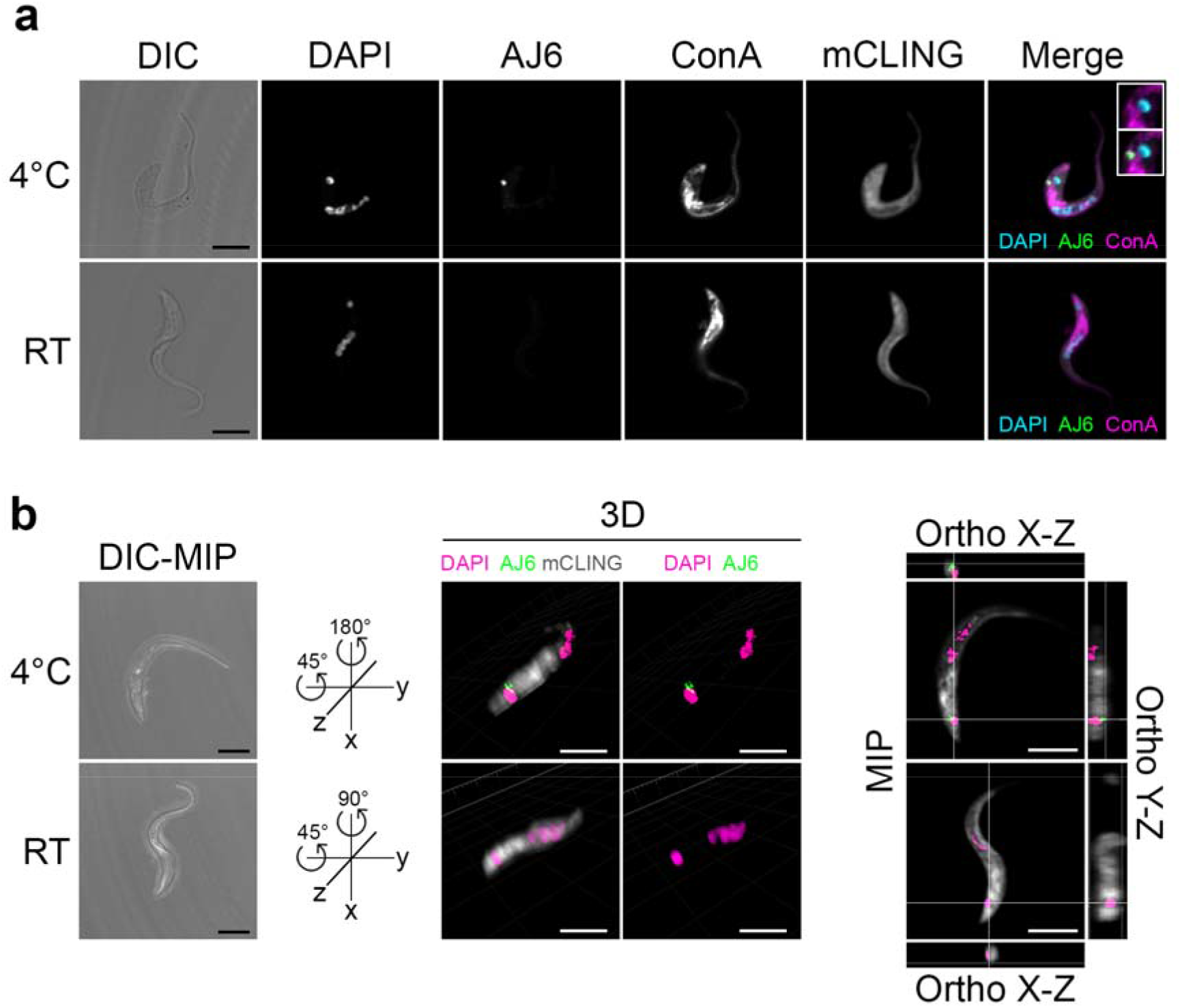
Live immunostaining of VIVβ8 antigen in native cells. **(A)** Live T. vivax bloodstream forms were immunostained either at 4°C to halt the secretory pathway or at room temperature (RT) to preserve it active. DIC; DAPI; VIVβ8 (secondary AF555-conjugated antibody); concanavalin A (ConA) FITC-conjugated lectin ER counterstain; mCLING unspecific staining and merged channels. Close-ups in merge channel show the flagellar pocket (end of ER next to kDNA) without (top) or with (bottom) VIVβ8 green signal. Scale bars; 5 μm. **(B)** 3D localization of VIVβ8 in live cells. Representative cells immunostained at 4°C or RT were 3D reconstructed in DIC-maximum intensity projection (DIC-MIP, left) and rotated to equivalent positions in the space to display VIVβ8 next to the kDNA only in 4°C cells (middle). MIP fluorescence projections (right) and orthogonal views. Scale bars; 5 μm.

Thus, VIVβ8 is certainly expressed on the plasma membrane, but there is a disparity between immunostaining of PFA-fixed cells, which implies staining of the whole cell surface, and of live cells, which indicates that VIVβ8 localises to the flagellar pocket only. This can be explained by epitope availability, if VIVβ8 is distributed across the whole cell membrane but obscured from antibody binding in its native state, perhaps by other proteins in the surface glycocalyx. However, PFA fixation exposes the epitopes, revealing its full distribution.

Finally, we also observed anti-VIVβ8 staining on red blood cells after PFA fixation of *T. vivax*-infected mouse blood. The stain localises in the centre of the bi-concave cell area, as shown by confocal microscopy (S4 Fig). It is unclear whether VIVβ8 is secreted actively by *T. vivax* to target the host erythrocytes, or if it is transferred accidentally after parasite cell lysis.

### Immunization with VIVβ8 produces a balanced antibody response

Having established the cell surface location of VIVβ8, we examined the potential of this protein family for vaccination. Initially, to establish a robust seroconversion, we inoculated BALB/c mice with our four recombinant vivaxin proteins in combination with multiple adjuvants and measured serum IgG1 and IgG2a antibody titres by indirect ELISA. Independently of adjuvant or antigen, seroconversion increased upon booster immunization indicating these antigens were immunogenic in mouse.

Antibody titres showed a significant increase (p<0.001) in both IgG1 and IgG2a-specific antibody compared with the pre-immune sera consistent response for all antigens regardless the adjuvant used (S5 Fig). However, adjuvant choice has a significant effect on antibody titres. Quil-A produced significantly higher IgG2a titres than either Montanide or alum when applied with all antigens. Overall, mice vaccinated with alum and Montanide elicited higher titres of IgG1 than IgG2a suggesting a strong Th2-type immune response, while Quil-A came closest to producing an equal ratio of isotype titres (ratio = 1.03), which indicates a mixed Th1/2-type response.

To compare the antibody responses to immunization with those observed for natural and experimental infections, we measured IgG1 and IgG2a titres in livestock serum seropositive for T. vivax (see above). Naturally-infected cattle from Cameroon and Kenya displayed significantly higher IgG1-specific titres than seronegative UK cattle (p < 0.05) for all four vivaxin antigens (S6 Fig). Experimentally infected cattle from Brazil showed a similar pattern to natural infections with a higher IgG1 than IgG2 antibody levels. Conversely, in most cases, anti-IgG2a responses for each antigen were not significantly greater than the negative control. These results indicate that, while vivaxin is strongly immunogenic in natural infections, it elicits a largely Th2-type response, similar that produced by immunization with Montanide or alum, but that immunization with Quil-A using any of the recombinant vivaxin antigens can produce a more balanced effect.

Cytokine expression provides further evidence for the type of immune response elicited by immunization. The concentrations of four cytokines (TNF-α, IFN-γ, IL-10 and IL-4) were measured in ex vivo mouse splenocyte cultures after stimulation with each vivaxin antigen, co-administrated with one of three adjuvants. All cytokines were undetectable in splenocytes cultured in media only, but after stimulation with an antigen, cytokine concentration increased significantly (p <0.0001; S7 Fig). Immunization with each antigen, regardless of adjuvant, produced high TNF-α concentration, with no significant differences between antigens (p > 0.05). IFN-γ concentration was greater in all animals immunized with Quil-A, which produced a similar response to the positive control group stimulated with ConA. The expression of IL-10 was also dependent on the adjuvant; it was significantly greater when antigens were co-administered with Quil-A (p <0.001 and p <0.0001 for all cases). IL-4 displayed the lowest expression levels of all; the only appreciate difference being for VIVβ20 co-administered with Quil-A compared to all other antigens (P < 0.001 for all cases). These results further indicate that immunization with vivaxin combined with Quil-A produces a balanced Th1-Th2 response.

### Vaccination with VIVβ8 with delays parasite proliferation

To evaluate the efficacy of vaccination, four mouse cohorts were vaccinated with a single vivaxin antigen and Quil-A respectively and were challenged with T. vivax bloodstream-forms (Fig 6A); parasitaemia was monitored by bioluminescent assay. In all cases, bioluminescence increased over the course of infection (Fig 6B). Before 5dpi, all vaccinated mice showed low parasitaemia levels similar to control groups, and showed no clinical symptoms. At 6dpi, the VIVβ8 cohort had the lowest parasitaemia with a mean of 2.45×108 p/s, while the other cohorts showed an average luminescence of 2.8×108 p/s. At 8dpi, when luminescence was greatest, the VIVβ20 cohort showed the highest parasitaemia of all groups, significantly greater than VIVβ11 (p=0.008) and VIVβ8 cohorts (p=0.002). By 9 dpi, however, all animals were sacrificed having developed clinical disease.

**Figure 6.**
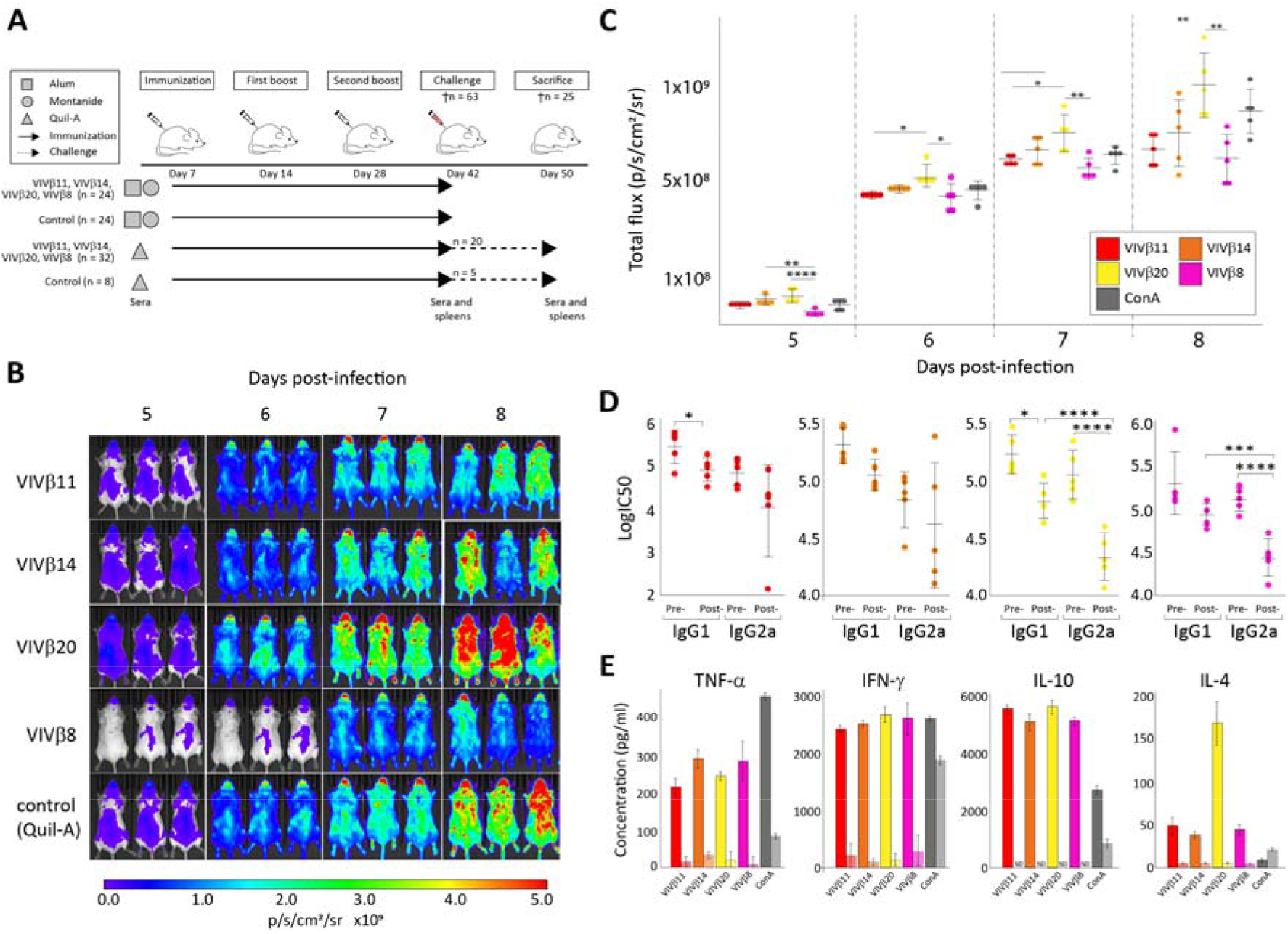
Vaccination and T. vivax challenge in a murine model. **(A)** Schedule of vaccine immunization and challenge. BALB/c mice were immunized every two weeks with each recombinant protein in conjunction with one of alum, Montanide or Quil-A adjuvants, while the control groups received adjuvants only. Animals were euthanized at day 42 (n=63) to assess the response to immunization, except for five mice from each vaccinated and control Quil-A-based groups (n=25) which were challenged with bloodstream-form T. vivax for a further eight days. **(B)** Vaccine protection against challenge with bioluminescent T. vivax in BALB/c mice. In vivo imaging of immunized mice with each of four vivaxin antigens co-administrated with Quil-A (n=5/group). Daily bioluminescent images were collected from 5-8dpi. **(C)** Parasite burden, measured as luminescent values (total flux in photons per second) of luciferase-expressing T. vivax in challenged mice at 8 DPI. **(D-E)** Cellular and humoral response before and after challenge with T. vivax in mice vaccinated with four antigens co-administered with Quil-A (n=8). Cytokine production by splenocytes stimulated in vitro (D) and isotype IgG profiling (E) were analyzed to determine the type of immune response elicited by each antigen.

At the end of the experiment, parasite luminescence in control and vaccinated animals was not statistically different overall (p > 0.05). However, this summary belies notable variation within the VIVβ8 cohort. Three of five VIVβ8 -vaccinated mice showed a delayed onset of parasite proliferation (Fig 6C), and a significant reduction in parasitaemia at 8dpi (p=0.045). Bioluminescence at 8dpi in the partially protected mice was 3.38×108 p/s compared to 7.17×108 p/s in the two unprotected VIVβ8 mice. This indicates that VIVβ8 adjuvanted with Quil-A inhibited parasite proflieration in some animals. We repeated the VIVβ8+Quil-A challenge using a larger cohort (n=15) and a greater amount of antigen (50μg) and this inhibited the onset of acute infection as before, with the bioluminescence from vaccinated animals being significantly lower than the control group at 6dpi (p =0.016; data not shown). However, there was no beneficial effect by 9dpi, with mice from vaccinated and control groups displaying a mean of 1.45×109 and 1.60×109 p/s, respectively.

After challenge, animals vaccinated with VIVβ14 showed a non-significant reduction in both IgG isotypes, while there was a significant reduction in the IgG1 titration in VIVβ11 and VIVβ20 vaccinated mice (p<0.01). IgG2a antigen-specific antibody levels also decreased significantly after challenge against VIVβ20 (p <0.0001) and VIVβ8 (p =0.0004; Fig 6D). Cytokine levels also displayed pronounced changes after challenge (Fig 6E), irrespective of the antigen involved. IL-10 expression became non-detectable after 8 dpi when compared with pre-vaccination levels (p<0.001). IL-4 concentrations also decreased significantly after challenge (p <0.0001). TNF-α and IFN-γ average concentrations against each antigen were reduced significantly (p <0.0001), representing a reduction of 95% and 92.8% respectively. Unstimulated cells from the adjuvant-only control group showed high cytokines levels indicating that Quil-A alone is able to stimulate their production.

Overall, all four vivaxin antigens were immunogenic, although they differed in the precise balance of immune response elicited, but none was able to protect against *T. vivax* infection in mouse. Only antigen-4, encoded by viv-*β*8, produced a balanced Th1-Th2 immune response after immunization with Quil-A and went on to have a significant negative effect on parasite burden. While encouraging, this effect was not observed in all replicates, and the balanced immune response was diminished after challenge with the decline of IgG2 titres relative to IgG1.

## Discussion

We examined the antibody responses of naturally and experimentally infected hosts to diverse TvCSP to identify immunogens that could become the basis for a *T. vivax* vaccine. Immunoprofiling of host serum showed that one particular family (Fam34), now known as vivaxin, is consistently the most immunogenic protein among the set we tested. Vivaxin is a large gene family encoding transmembrane glycoproteins with a conserved primary structure but diverse expression profiles and population genetic dynamics. Thus, not all vivaxin genes are equally good candidate antigens; we focused on one gene with minimal polymorphism (viv-*β*8) that was among the most immunogenic and highly expressed, and confirmed its expression across the cell body and flagellar membranes.

While VIVβ8 elicits a strong, balanced immune response with Quil-A and significantly reduces parasite burden in some mice (through delayed parasite proliferation), animals were not protected from fatal disease. In fact, the experiment followed a familiar pattern, with reduced antibody titres and pro-inflammatory cytokine concentrations after challenge. Reduction of IgG1 and IgG2a antibody titres before and after challenge was observed previously in vaccinated cattle challenged with both *T. vivax* [48] and *T. congolense* [49], as well as vaccinated mice challenged with *T. brucei* [21]. A possible explanation for this event could the reduction in number of splenic B-cells which is associated with a defect in B-cell development [50]. Another possible explanation could be the formation of immune complex of specific antibodies with the parasite antigens making this unable to measure IgGs solely in circulation [51-52]. Reduction of antigen-stimulated cytokine levels observed after 8 dpi is not conducive to parasite control, since IFN-γ is associated with resistance to African trypanosomes [53-55] and TNF-α was shown to be essential to controlling *T. vivax* infection in mice [56]. These changes in both the IgG1/IgG2a ratio and the cytokine profiles after challenge indicate a transition from a Th1-type to Th2-type response, which is a typical feature of uncontrolled infections in trypanosomatids [57-61] and is observed in naturally-infected cattle [62].

Thus, after vaccination with VIVβ8, infection took its normal course. It could be that either vivaxin epitopes are hidden in situ or that bound vivaxin is removed via endocytosis under physiological conditions. Immunofluorescent microscopy of fixed trypomastigotes using both purified recombinant antibodies (Fig.3) and post-immune serum (Fig.4) indicated that VIVβ8 is located across the entire cell surface, and so, a uniform component of the glycocalyx alongside the VSG. However, when antibodies were applied to live parasites they fail to bind, except when the cells are cooled, and then only to the flagellar pocket (Fig.5). Typically, we might suggest that the VSG is responsible for obscuring vivaxin from antibodies everywhere except the flagellar pocket, but most α-vivaxin predicted proteins are 1.5-2 times larger than a typical *T. vivax* VSG (□ 450 amino acids), making this problematic.

If vivaxin epitopes are inaccessible in vivo, we must explain the strong and consistent serological response of both naturally and experimentally-infected animals to multiple vivaxin proteins. The strength of the serological response perhaps reflects the abundance and conservation of vivaxin, since we linked the most abundant and least polymorphic transcripts to the strongest immunogens. However, vivaxin may be secreted and eliciting antibodies after cleavage of the extracellular domain from the cell surface, or else, the antibody response may be directed primarily at dead and lysed parasites. Such responses would not affect live, circulating parasites if the protein remained concealed on their surface. Certainly, VIVβ8 protein attaches to erythrocytes in vivo and, although the physiological effects of this is unknown, it might serve to dilute the effective antibody response.

Although VIVβ8 may not elicit protective immunity, our phylogenetic analysis reveals that two antigens that were effective in a previous study, IFX and V31 [23], belong to the vivaxin gene family. We now realize the IFX (VIVβ1) is also among the most strongly expressed and structurally conserved vivaxin proteins (although not as much as VIVβ8). V31 (VIVα18) had a partially protective effect in mice, but is much more polymorphic [23]. Curiously, viv-*β*1 (encoding IFX) adopts a unique position within the phylogeny, as the sister lineage to all other *β*-vivaxin. This topology is highly robust but nonetheless odd, because viv-*β*1 is much longer than other *β*-vivaxin and noticeably divergent (note the length of the viv-*β*1 branch). It is tempting to speculate from the strong conservation and divergent structure of VIVβ1 that this protein performs a distinct, non-redundant function among vivaxin proteins, a function that evidently exposes it to antibodies unlike many of its paralogs. Note that while VIVβ8 localizes across the cell body and flagellum, VIVβ1 was restricted to regions of the flagellar membrane [23]. Thus, among the 124 (and likely more) vivaxin paralogs there is great potential for reliably immunogenic and protective antigens; yet this study reveals substantial variability in structure and antigenic properties, even among closely related gene copies, such that not all vivaxin proteins will make good antigens.

Besides its potential for subunit vaccines, the discovery of vivaxin has implications for host-parasite interactions. The protein architecture of the *T. vivax* cell surface is not well characterised, partly because attention is more typically focused on the human pathogen T. brucei, but also because there are few research tools (e.g., in vitro cell culture, reverse genetics, mouse infection model) developed for T. vivax [63]. Yet, recent results and historical anecdote suggest that the T. vivax cell surface is quite different to the uniform and pervasive VSG monolayer of *T. brucei*. Vickerman considered the *T. vivax* surface coat to be less dense than other species [64-65]. The T. vivax genome contains hundreds of species-specific and non-VSG genes [24,66]. Greif et al. (2013) showed that only 57% of transcripts during *T. vivax* mouse infections encoded VSG (compared to 98% of T. brucei bloodstream-stage transcripts) and that the remainder belonged largely to T. vivax-specific genes [44]. This study shows that vivaxin must be a major contributor to this difference between T. vivax and T. brucei surfaces.

It follows that, with a different cell surface architecture, *T. vivax* may interact with hosts in a different way, dependent on what the function(s) of vivaxin might be. Other trypanosome surface proteins are variant antigens (e.g. VSG [67]), immunomodulators in other ways (e.g. trans-sialidases [68]), scavenge nutrients (e.g. transferrin and HpHb receptors [69-70]) or sense the host environment (e.g. adenylate cyclases [71-72]). Various molecular evolutionary aspects, (i.e. strong purifying selection, low polymorphism, maintenance of gene orthology), as well as the absence of monoallelic expression and abundant glycosylation sites, show that vivaxin are not variant antigens. However, other functions in immunomodulation or environmental sensing are plausible. The secretion (active or otherwise) of VIVβ8 and its adhesion to erythrocytes (S3 Fig) also suggests that vivaxin could potentially facilitate cytoadhesion, leading to parasite sequestration in tissue capillaries as an immune evasion strategy.

Perhaps the only aspect of vivaxin function we can predict presently is that it will be multifarious. Differences in length among subfamilies will translate into distinct tertiary protein structures, while consistent differences in expression profile suggest that some vivaxin genes are ‘major forms’, while other paralogs appear to be non-functional, and a few may be expressed beyond the bloodstream-stage. Population genetics show that vivaxin genes evolve under a range of selective conditions, from strongly negative (i.e. functionally essential and non-redundant), to neutral (i.e. redundant), and positive (engaged in antagonistic host interactions?). Coupled with the evolutionary stability of these features, (that is, individual vivaxin genes are found in orthology across *T. vivax* strains rather than recombining or being gained and lost frequently), this is evidence for functional differentiation and non-redundancy within the gene family.

Vivaxin represents a major component of the *T. vivax* surface coat, quite distinct from VSG, and includes proven vaccine targets, and many more potential targets. The molecular evolution of vivaxin implies that the paralogous gene copies lack the dynamic variability and redundancy of variant antigens, but instead perform multiple distinct and essential functions. The discovery of this highly immunogenic and abundant protein family has important implications for how we understand AAT caused by *T. vivax*. It challenges the adequacy of *T. brucei* as a model for AAT, given the different qualities of their surface architectures, while posing new therapeutic opportunities and new questions about the roles vivaxin has in host interaction, immune modulation and disease.

## Acknowledgements

This work was supported by BBSRC research grants (BB/S001980/1 and BB/R021139/1) and a pump-priming grant from the International Veterinary Vaccinology Network to APJ. ARR was supported by a doctoral studentship by FONDECYT-CONCYTEC, the National Council of Science, Technology and Innovation from Peru (grant contract number 001-2016-FONDECYT). We thank Ben Makepeace (University of Liverpool) and Sara Silva Pereira (Instituto de Medicina Molecular, Lisbon) for kind donations of bovine serum samples.

## Supporting Information

**S1 Fig. Peptide microarray slide design**. The diagram shows the 600 spots of the microarray, with each cell corresponding to two peptides printed in duplicate (scale at edge). Each spot contains a 15-mer peptide belonging to one of 62 *Trypanosoma vivax proteins*, printed with a 14 amino acid overlap, or a control peptide. The cells are shaded to identify the *T. vivax* cell surface phylome (TvCSP) to which each non-control peptide belongs [24]. Twenty proteins do not belong to multi-copy families (‘Single-copy’), but are still predicted to have cell surface expression.

**S2 Fig. Predicted secondary protein structures for six vivaxin genes**. The six genes include those four encoding antigens 1-4 identified in this study and expressed in recombinant form (viv-*β*11, viv-*β*14, viv-*β*20 and viv-*β*8), as well as two others encoding candidate antigens from another study (viv-*β*1 and viv-*α*18; [23]) for comparison. Protein secondary structures were inferred from amino acid sequences using PredictProtein [41]: alpha helices (red), transmembrane helix (purple), disordered region (green). The solvent accessibility of each position is also indicated: accessible (blue) and buried (yellow). N- and O-linked glycosylation sites were predicted using ModPred [42] and are indicated by red and orange arrows respectively. The position of linear b-cell epitopes inferred from the TvCSP peptide microarray are indicated by grey bars at the bottom of each diagram (the range of positions in the amino acid sequence is given).

**S3 Fig. Recombinant expression of four vivaxin proteins**. A) Normalization of viv-*β*11 protein using two-fold serial dilutions. B) Normalization of VIVβ14, VIVβ20 and VIVβ8 proteins. C) Western blot analysis confirming protein expression. One microgram of each antigen were separated on a 12% acrylamide gel and transferred to a nitrocellulose membrane and detected using an anti-biotin secondary antibody HRP (right). M: molecular weight marker.

**S4 Fig. Cellular localization of VIVβ8 antigen on the surface of murine erythrocytes**. (A) Localization of VIVβ8 and the unspecific surface counterstain mCLING in red blood cells (RBC) from *T. vivax*-infected mice. Representative images of RBC stained with either pre-immune or post-immune rabbit polyclonal antisera. Middle row shows the major localization pattern of VIVβ8 in RBC; protein accumulates in the central concave surface. Bottom row shows an example of leaking RBC. Differential increased contrast (DIC); DAPI DNA counterstain; VIVβ8 (secondary antibody AF555-conjugated) and merged channels. Scale bars; 5 μm. (B) 3D z-stack reconstructions of mouse erythrocyte cells and corresponding orthogonal (X-Z and X-Y) views from the stacks. Orthogonal views note the VIVβ8 signal originates in the inner concave cytoplasm. Scale bars; 5 μm.

**S5 Fig. Antibody titres after immunization**. Both IgG1 and IgG2a-specific antibody compared with the pre-immune sera consistent response for all antigens regardless the adjuvant used. However, adjuvant choice has a significant effect on antibody titres. Montanide produced significantly higher IgG1 levels than either alum or Quil-A when applied with VIVβ11, VIVβ14 and VIVβ20, but, there was no difference in IgG1 titre between adjuvants when VIVβ8 was used. In contrast, Quil-A produced significantly higher IgG2a titres than either Montanide or alum when applied with all antigens.

**S6 Fig. IgG1 and IgG2a titres in livestock serum seropositive for *T. vivax***. Naturally infected cattle from Cameroon and Kenya displayed significantly higher IgG1-specific titres than seronegative UK cattle (p < 0.05) for all four vivaxin antigens.

**S7 Fig. Cytokine expression after immunization compared for different adjuvants**. The concentrations of four cytokines (TNF-α, IFN-γ, IL-10 and IL-4) were measured in ex vivo mouse splenocyte cultures after stimulation with each vivaxin antigen, co-administrated with one of three adjuvants.

