## Supplementary Figure 1 for "*Vivaxin* genes encode highly immunogenic non-variant antigens unique to the *Trypanosoma vivax* cell-surface"

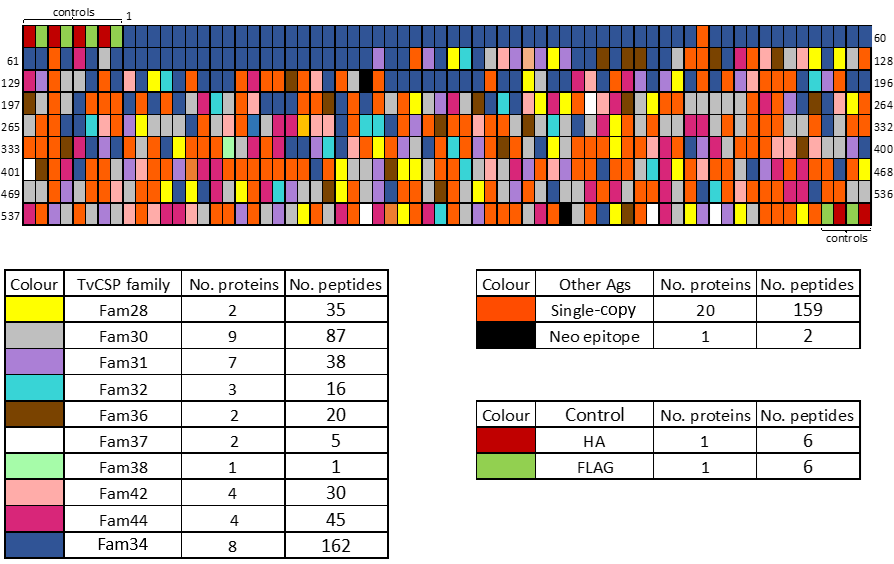

**Supplementary Figure 1. Peptide microarray slide design**. The diagram shows the 600 spots of the microarray, with each cell corresponding to two peptides printed in duplicate (scale at edge). Each spot contains a 15-mer peptide belonging to one of 62 *Trypanosoma vivax* proteins, printed with a 14 amino acid overlap, or a control peptide. The cells are shaded to identify the *T. vivax* cell surface phylome (TvCSP) to which each non-control peptide belongs [24]. Twenty proteins do not belong to multi-copy families (‘Single-copy’), but are still predicted to have cell surface expression.
