## Supplementary Figure 2 for "*Vivaxin* genes encode highly immunogenic non-variant antigens unique to the *Trypanosoma vivax* cell-surface"

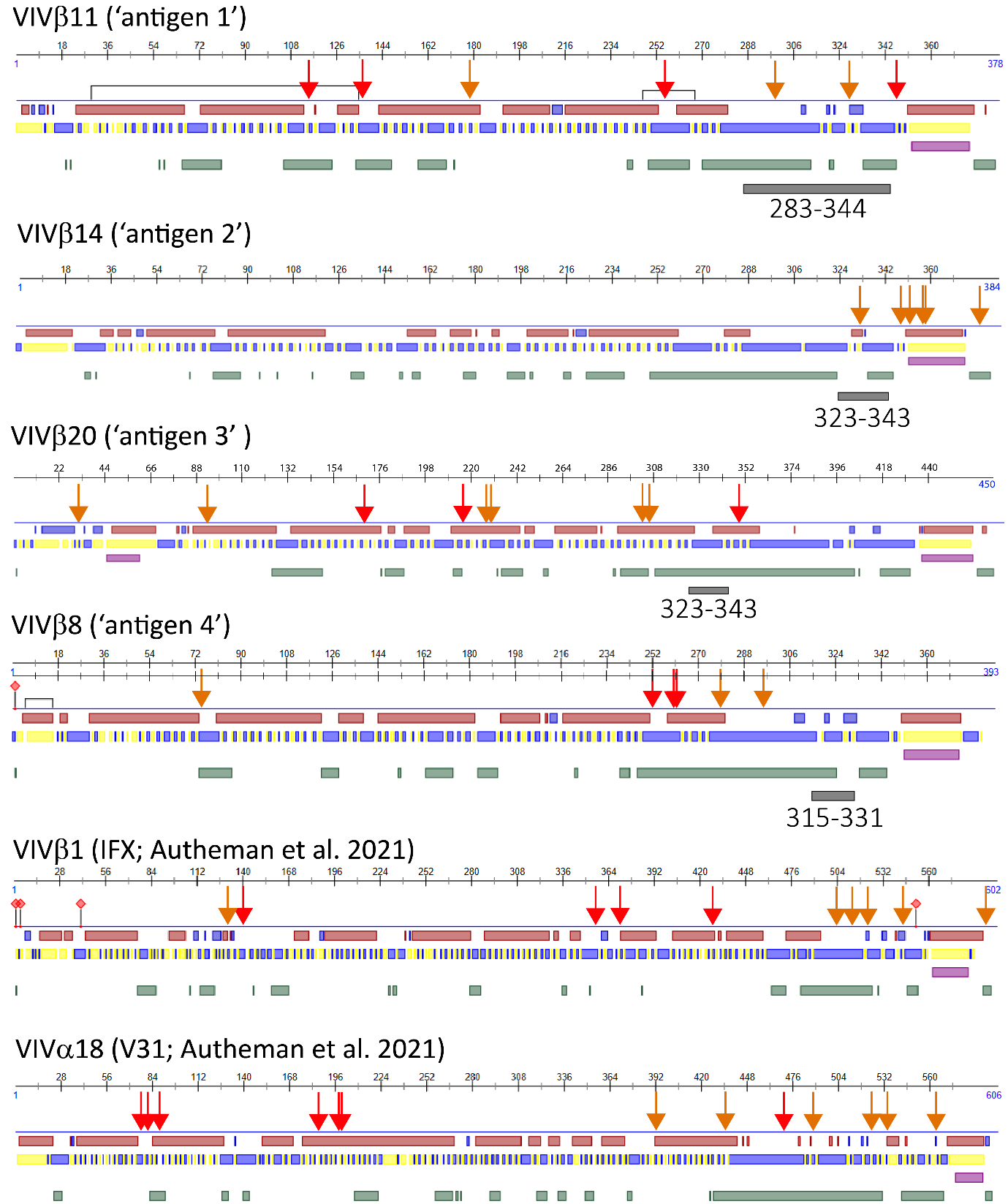

**Supplementary Figure 2. Predicted secondary protein structures for six vivaxin genes.** The six genes include those encoding four antigens identified in this study and expressed in recombinant form (*viv-β11, viv-β14*, *viv-β20* and *viv-β8*), as well as two others encoding candidate antigens from another study (*viv-β1* and *viv-α18*; Autheman et al. 2021) for comparison. Protein secondary structures were inferred from amino acid sequences using PredictProtein (Yachdav et al. 2014): alpha helices (red), transmembrane helix (purple), disordered region (green). The solvent accessibility of each position is also indicated: accessible (blue) and buried (yellow). N- and O-linked glycosylation sites were predicted using ModPred (Pejaver et al. 2014) and are indicated by red and orange arrows respectively. The position of linear b-cell epitopes inferred from the TvCSP peptide microarray are indicated by grey bars at the bottom of each diagram (the range of positions in the amino acid sequence is given).
