## Supplementary Figure 3 for "*Vivaxin* genes encode highly immunogenic non-variant antigens unique to the *Trypanosoma vivax* cell-surface"

Antigen-2

Antigen-3

Antigen-4

**Antigen:**

**M	1	2	3	4**

Antigen-1

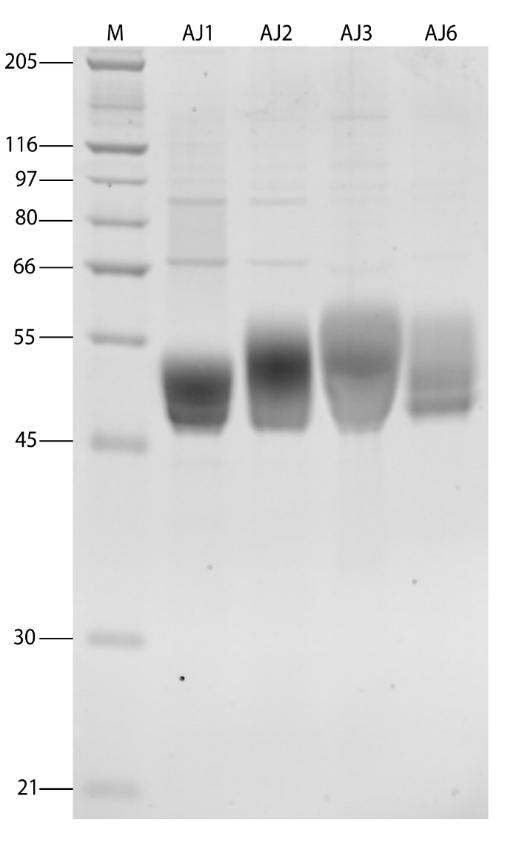

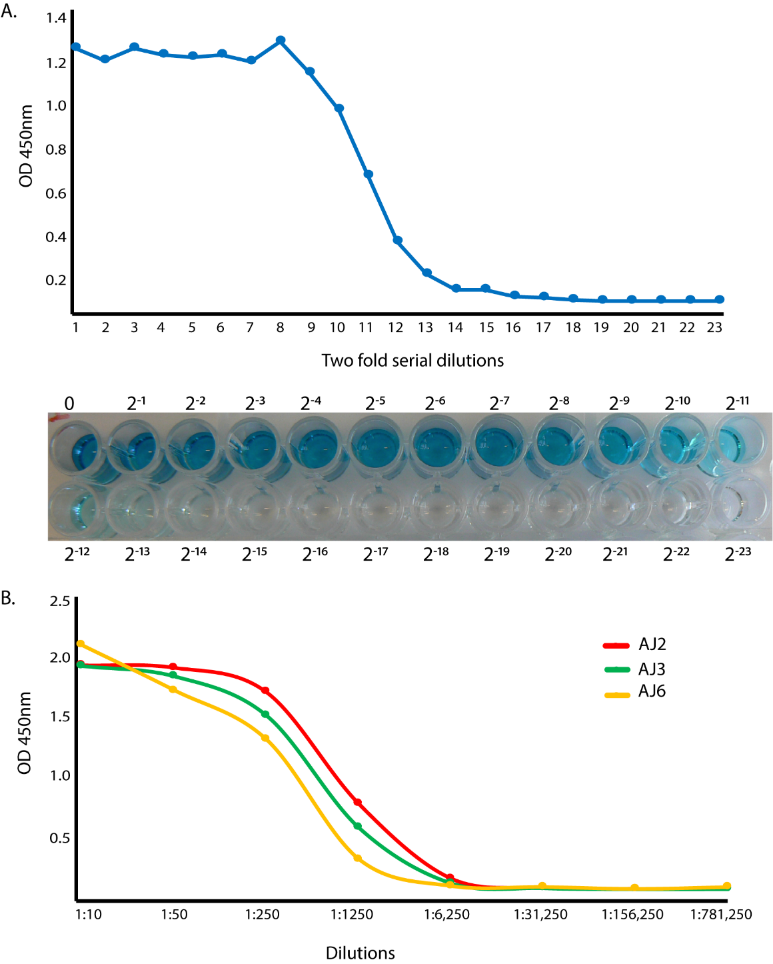

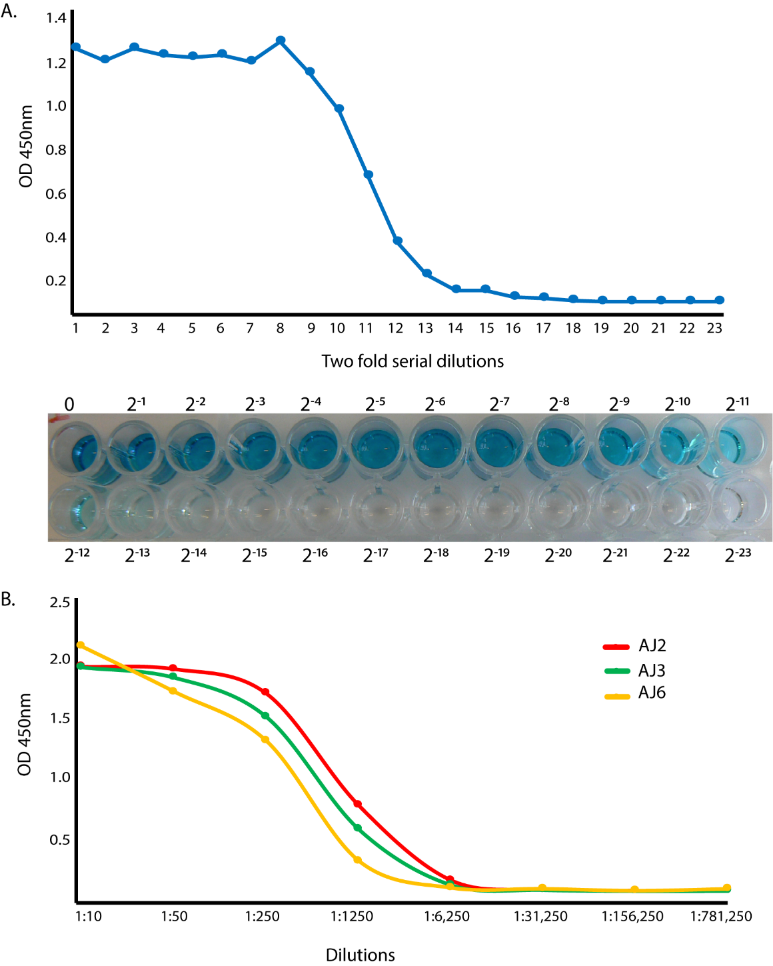

**Supplementary Figure 3. Recombinant expression of four vivaxin proteins.** A) Normalization of ‘antigen 1’ (VIVβ11) protein using two-fold serial dilutions. B) Normalization of ‘antigen 2’ (VIVβ14), ‘antigen 3’ (VIVβ20) and ‘antigen 4’ (VIVβ8) proteins. C) Western blot analysis confirming protein expression. One microgram of each antigen were separated on a 12% acrylamide gel and transferred to a nitrocellulose membrane and detected using an anti-biotin secondary antibody HRP (right). M: molecular weight marker.
