## Supplementary Figure 4 for "*Vivaxin* genes encode highly immunogenic non-variant antigens unique to the *Trypanosoma vivax* cell-surface"

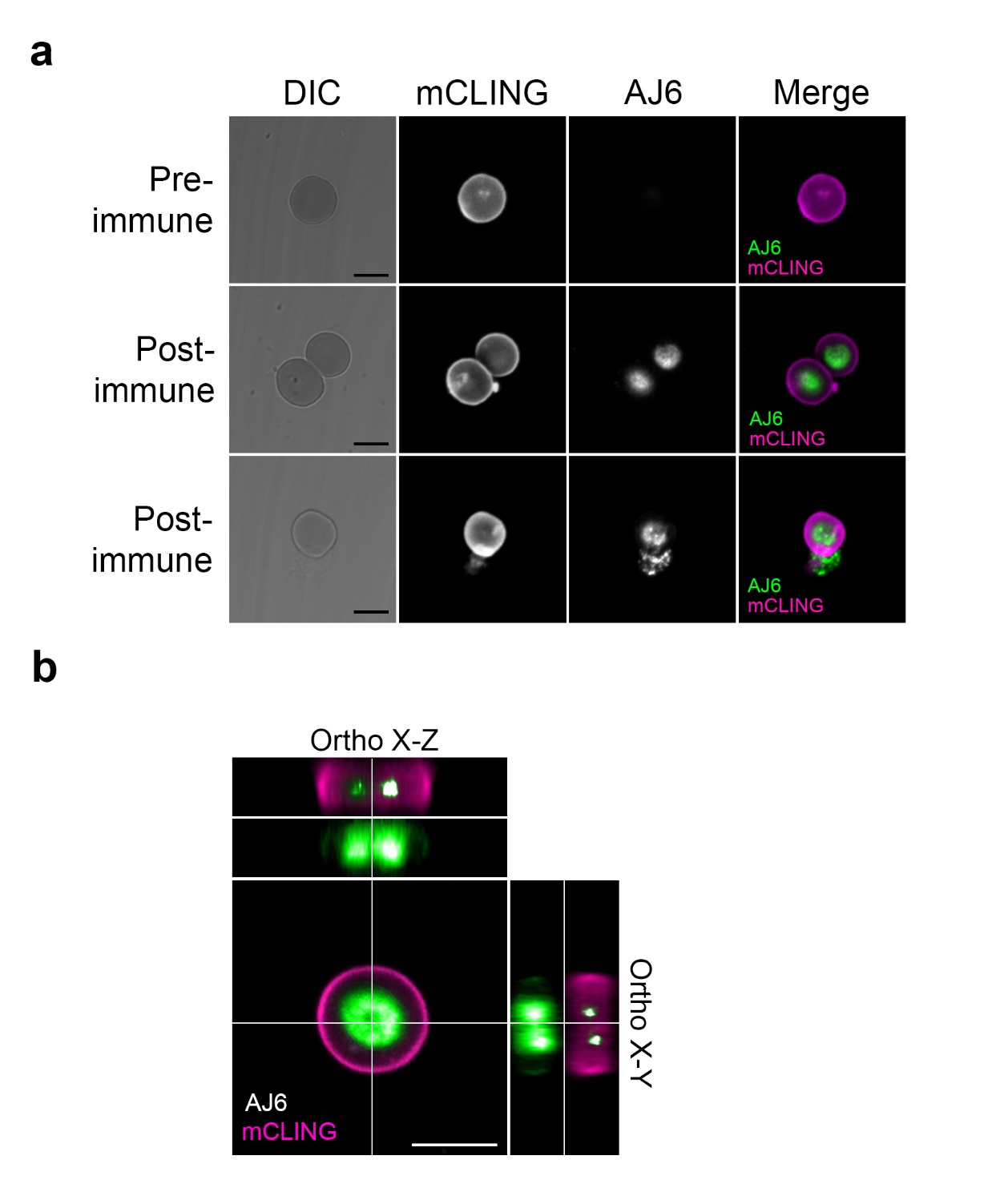

**Supplementary Figure 4**. **Cellular localization of VIVβ8 antigen on the surface of murine erythrocytes**. **(A)** Localization of VIVβ8 and the unspecific surface counterstain mCLING in red blood cells (RBC) from *T. vivax*-infected mice. Representative images of RBC stained with either pre-immune or post-immune rabbit polyclonal antisera. Middle row shows the major localization pattern of VIVβ8 in RBC; protein accumulates in the central concave surface. Bottom row shows an example of leaking RBC. Differential increased contrast (DIC); DAPI DNA counterstain; VIVβ8 (secondary antibody AF555-conjugated) and merged channels. Scale bars; 5 μm. **(B)** 3D z-stack reconstructions of mouse erythrocyte cells and corresponding orthogonal (X-Z and X-Y) views from the stacks. Orthogonal views note the VIVβ8 signal origins in the inner concave cytoplasm. Scale bars; 5 μm.
