## Supplementary Figure 5 for "*Vivaxin* genes encode highly immunogenic non-variant antigens unique to the *Trypanosoma vivax* cell-surface"

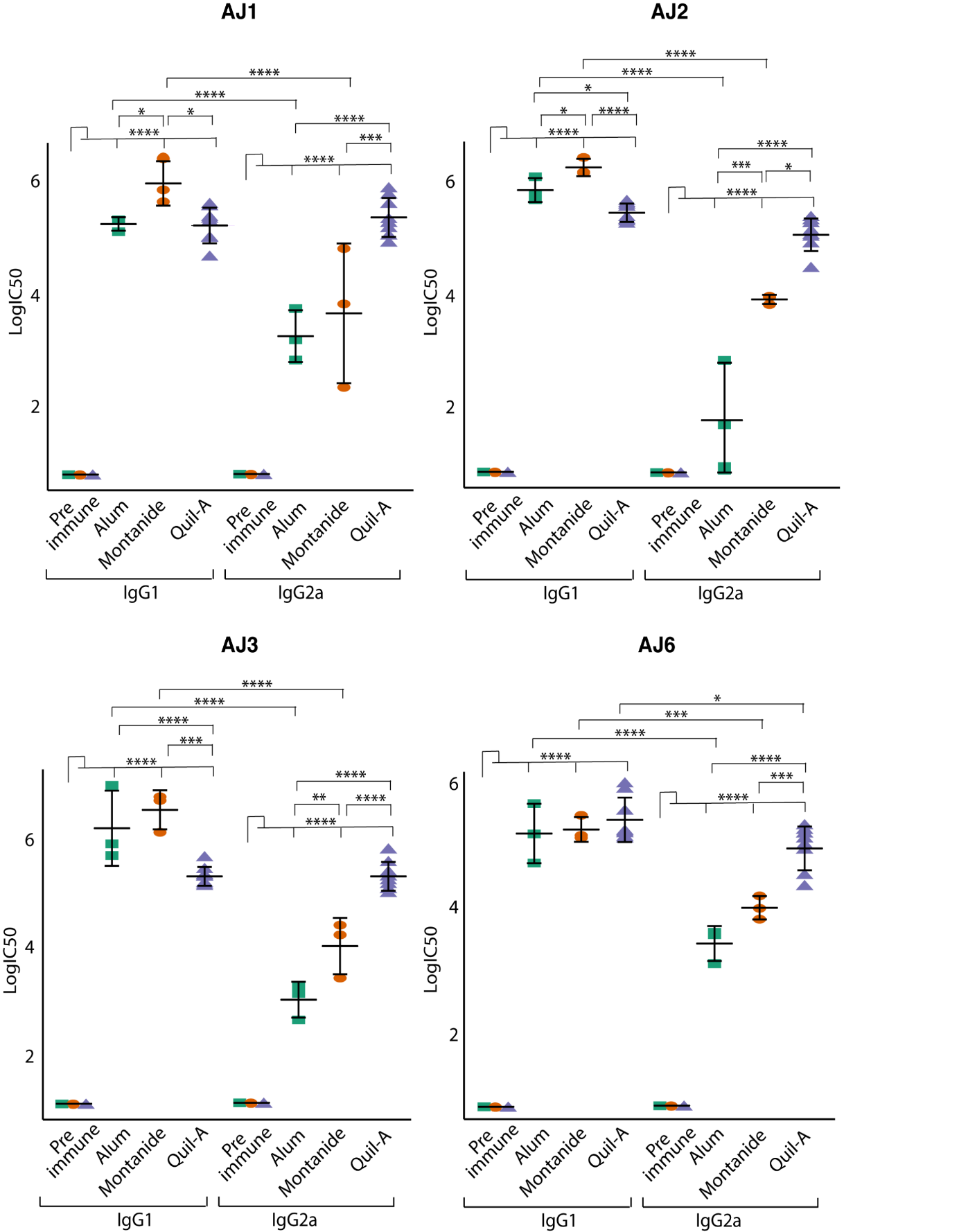

**A.**

**B.**

**C.**

**D.**

**Antigen-1**

**Antigen-2**

**Antigen-3**

**Antigen-4**

**Supplementary Figure 5**. **Murine immune response to immunization.** Titration of *T. vivax* antigens-specific IgG1 and IgG2a antibody response in BALB/c mice before and after immunization with one of antigens 1-4 (n=4 for each antigen, except n=8 for Quil-A pre and post vaccination). IgG1 and IgG2a specific antibody titres were measured by ELISA using two-fold serially diluted sera. Data are represented as the average of triplicates.
