## Supplementary Figure 6 for "*Vivaxin* genes encode highly immunogenic non-variant antigens unique to the *Trypanosoma vivax* cell-surface"

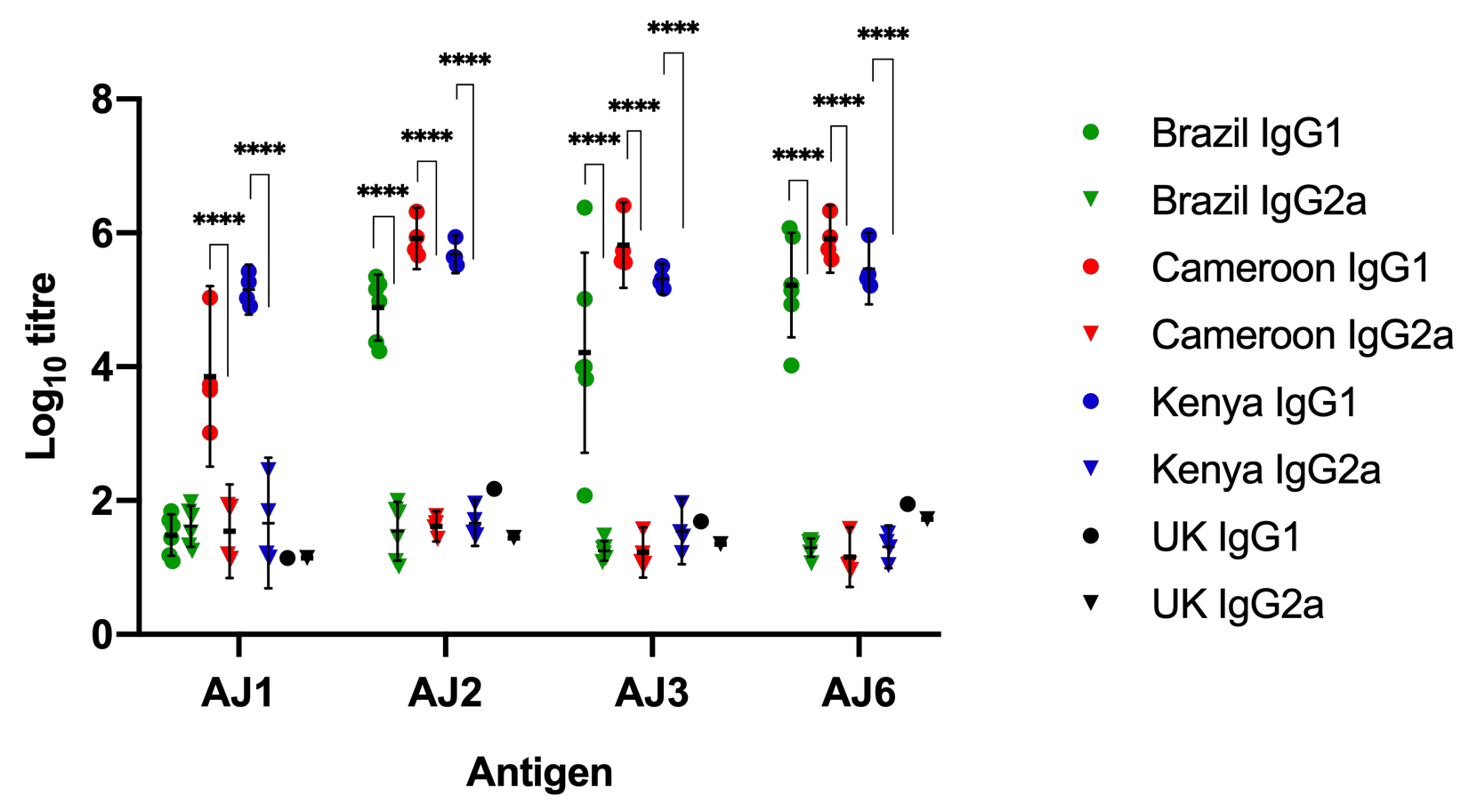

**1	2	3	4**

Antigen-1

**Supplementary Figure 6. Titre of IgG1 and IgG2a isotypes in infected cattle against antigens 1-4, measured by indirect ELISA**. IgG1 and IgG2a specific antibody titres were measured using two-fold serial dilutions in naturally infected (Cameroon and Kenya) and experimentally infected cattle (Brazil). Antibody levels were also measured in a group of UK cattle, which served as negative controls. IgG1 showed higher levels when compared to IgG2a for both natural and experimental infections with *T. vivax*. Each graph shows the antibody levels of individual serum, the geometric mean of each group, and the 95% confidence interval. ****, statistically significant at *P* < 0.0001.
