## Supplementary Figure 7 for "*Vivaxin* genes encode highly immunogenic non-variant antigens unique to the *Trypanosoma vivax* cell-surface"

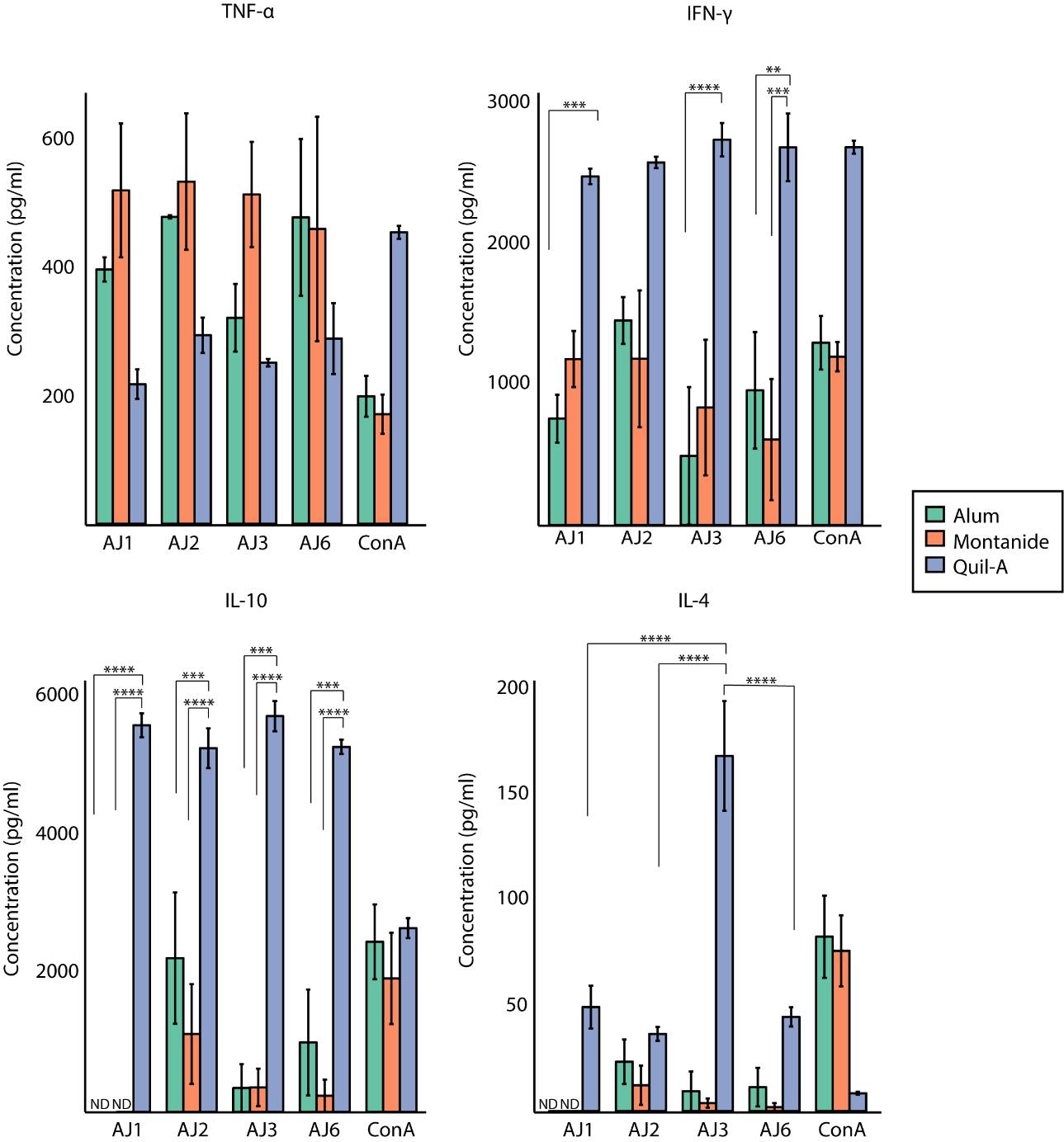

**1	2	3	4	ConA**

**1	2	3	4	ConA**

**1	2	3	4	ConA**

**1	2	3	4	ConA**

A.

B.

D.

C.

**Supplementary Figure 7.** **Cytokine responses of *in vitro* stimulated splenocytes from BALB/c mice vaccinated with one of antigens 1-4 prior to challenge***.* ND: non detectable. P-value < 0.05.
